# Structural basis for repurpose and design of nucleoside drugs for treating COVID-19

**DOI:** 10.1101/2020.11.01.363812

**Authors:** Wanchao Yin, Xiaodong Luan, Zhihai Li, Yuanchao Xie, Ziwei Zhou, Jia Liu, Minqi Gao, Xiaoxi Wang, Fulai Zhou, Qingxia Wang, Qingxing Wang, Dandan Shen, Yan Zhang, Guanghui Tian, Haji A. Aisa, Tianwen Hu, Daibao Wei, Yi Jiang, Gengfu Xiao, Hualiang Jiang, Leike Zhang, Xuekui Yu, Jingshan Shen, Shuyang Zhang, H. Eric Xu

**Affiliations:** The CAS Key Laboratory of Receptor Research, Shanghai Institute of Materia Medica, Chinese Academy of Sciences, Shanghai 201203, China; School of Medicine, Tsinghua University, Haidian District, Beijing, China; Department of Cardiology, Peking Union Medical College Hospital, Peking Union Medical College and Chinese Academy of Medical Sciences, Beijing, China; Tsinghua-Peking Center for Life Sciences, Tsinghua University, Beijing, China; Cryo-Electron Microscopy Research Center, Shanghai Institute of Materia Medica, Chinese Academy of Sciences, Shanghai 201203, China; University of Chinese Academy of Sciences, Beijing 100049, China; WuxiBiortus Biosciences Co. Ltd., 6 Dongsheng West Road, Jiangyin 214437, China; State Key Laboratory of Virology, Wuhan Institute of Virology, Center for Biosafety Mega-Science, Chinese Academy of Sciences, Wuhan, Hubei, 430071, P. R. China; Department of Biophysics, and Department of Pathology of Sir Run Run Shaw Hospital, Zhejiang University School of Medicine, Hangzhou 310058, China; Key Laboratory of Immunity and Inflammatory Diseases of Zhejiang Province, Hangzhou 310058, China; Vigonvita Life Science Co., Ltd., Suzhou 215123, China; Key Laboratory of Plant Resources and Chemistry in Arid Regions, Xinjiang Technical Institute of Physics and Chemistry, Chinese Academy of Sciences, South Beijing Road 40-1, Urumqi, Xinjiang 830011, P. R. China

## Abstract

SARS-CoV-2 has caused a global pandemic of COVID-19 that urgently needs an effective treatment. Nucleoside analog drugs including favipiravir have been repurposed for COVID-19 despite of unclear mechanism of their inhibition of the viral RNA polymerase (RdRp). Here we report the cryo-EM structures of the viral RdRp in complex with favipiravir and two other nucleoside inhibitor drugs ribavirin and penciclovir. Ribavirin and the ribosylated form of favipiravir share a similar ribose scaffold that is distinct from penciclovir. However, the structures reveal that all three inhibitors are covalently linked to the primer strand in a monophosphate form despite the different chemical scaffolds between favipiravir and penciclovir. Surprisingly, the base moieties of these inhibitors can form mismatched pairs with the template strand. Moreover, in view of the clinical disadvantages of remdesivir mainly associated with its prodrug form, we designed several orally-available remdesivir parent nucleoside derivatives, including VV16 that showed 5-fold more potent than remdesivir in inhibition of viral replication. Together, these results demonstrate an unexpected promiscuity of the viral RNA polymerase and provide a basis for repurpose and design of nucleotide analog drugs for COVID-19.

**One Sentence Summary:** Cryo-EM structures of the RNA polymerase of SARS-CoV-2 reveals the basis for repurposing of old nucleotide drugs to treat COVID-19.

---

Severe acute respiratory syndrome coronavirus 2 (SARS-CoV-2), which causes the pandemic of COVID-19, has continued to infect many people across the world (*1, 2*). SARS-CoV-2 is related to the highly pathogenic beta-coronaviruses that include SARS-CoV and Middle East respiratory syndrome coronavirus (MERS-CoV)(*3*). These viruses are positive-sense, single-stranded RNA viruses and their infections could lead to severe acute respiratory syndrome, which could result in loss of lung function and death. Among these related coronaviruses, SARS-CoV-2 has much higher activity to spread among humans, resulting in a global scale of health crisis, which still needs an effective treatment. Drug development usually requires a decade long of effort and it is unfit for current medical crisis. Repurpose of existing drugs that bypass regular preclinical and clinical research is of tremendous medical importance to combat the pandemics of COVID-19.

The viral RNA-dependent RNA polymerase (RdRp) is essential for viruses to replicate and has served as a major drug target for antiviral therapy. Numerous nucleotide analogs, including favipiravir(*1, 5*), ribavirin(*1, 7*), and penciclovir(*8*), have been developed to target the polymerase of different viruses such as influenza virus, hepatitis C virus (HCV), and Zika virus (ZIKV). The recent structure of the SARS-CoV-2 RdRp in complex with RNA substrate and nucleotide inhibitor remdesivir has revealed a conserved active site(*1, 10*), which is highly similar to the active site of the RdRp from several viruses, suggesting that nucleotide inhibitors might have broad antiviral activity. Indeed, favipiravir and ribavirin, which are well established drugs for influenza virus and hepatitis C virus (HCV), have been repurposed for the treatment of SARS-CoV-2(*11-14*). Remdesivir has been approved by FDA for treating COVID-19 and favipiravir has been approved in China, Russia, and India for COVID-19(*1, 16*). In addition, in this paper we also found that penciclovir, a nucleotide analog drug for herpesvirus, is an inhibitor of the SARS-CoV-2 RdRp, despite that penciclovir lacks a ribose scaffold like ribavirin. However, the lack of structural details of how these nucleotide analog drugs are recognized by RdRp has hampered the development of more potent and effective inhibitors to treat SARS-CoV-2 infections. In this paper, we report three structures of the SARS-CoV-2 RdRp in complex with RNA substrate and favipiravir, ribavirin, and penciclovir, respectively. Based on these structures and previous remdesivir-bound structures, we have designed several orally-available derivatives of remdesivir with potent antiviral activity. Together, these results reveal an unexpected promiscuity of the viral RNA polymerase and provide structural mechanisms for repurpose and design of nucleotide analog drugs for treating COVID-19.

The functional assembly of the SARS-CoV-2 RNA polymerase requires minimal three non-structural proteins, the RdRp subunit nsp12 with two accessary subunits nsp7 and nsp8(*1, 17*). Incubation of the purified nsp12-7-8 complex (fig. S1A-S1B) with a 30-base template, which had been annealed with a 20-base primer (Fig.1A), allowed the primer extension to the same length as the template in the presence of NTP (Fig.1B-1D). Interestingly, we also observed a product with one nucleotide longer than the template, indicating that RdRp may have a template-independent terminal nucleotide transferase activity (fig. S1C). It has been known that nucleotide analog drugs need to be converted into the active triphosphate form (fig. S2). Consistently, incubation of favipiravir, ribavirin, and penciclovir did not inhibit the primer extension by the RdRp (fig. S3). However, when favipiravir, ribavirin, and penciclovir were made with their corresponding triphosphate forms, they were able to inhibit the RdRp primer extension activity (Fig.1B-1D). The triphosphate form of favipiravir (FAV-TP), as expected, acted like an ATP analog and was able to inhibit the RdRp extension on a poly-U template (Fig.1B). Unexpectedly, FAV-TP was also able to inhibit the RdRp activity on poly-A, poly-G, and poly-C templates (Fig.1B), indicating that the base analog of favipiravir is able to form non-selective base pairing interactions with A, G, C, U bases. We also noticed that the SARS-CoV-2 RdRp had a relatively low processivity on a poly-G template when compared to poly-U and poly-A templates (Fig.1B-1D).

**Fig. 1.**
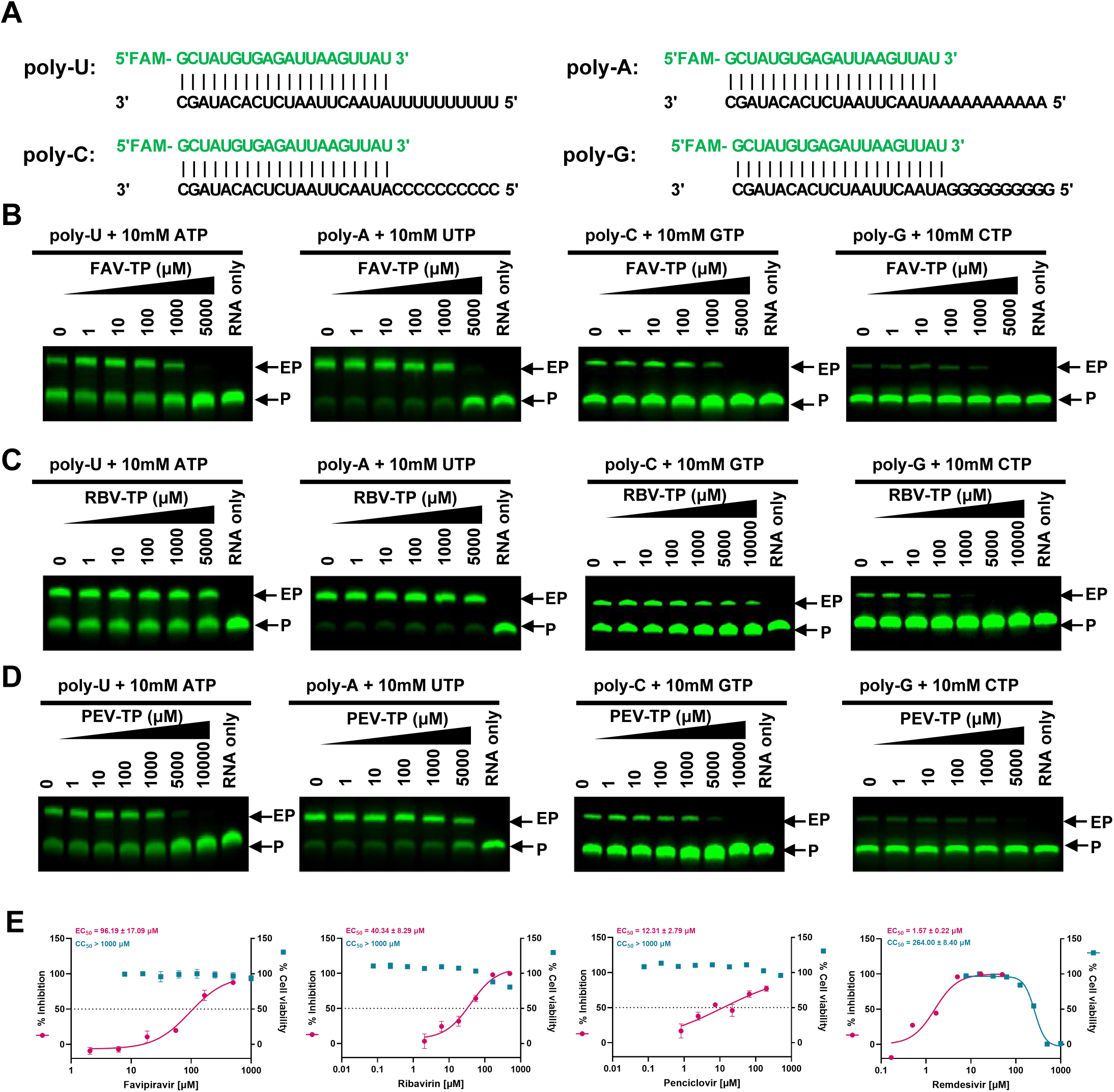
Specific Inhibition of RdRp by nucleotide drugs. A, Diagram of the 30-base template and 20-base primer duplex RNAs with a FAM at the 5’ of the primer used in the gel-based elongation assay. B-D, Inhibitions of RNA elongation with different substrates by FAV-TP (B), RBV-TP (C), and PEV-TP (D). The EP means the elongated product, while the P means the primer RNA strand. E, Inhibition of SARS-CoV-2 replication (EC_50_) and cellular toxicity (CC_50_) of remdesivir, favipiravir, ribavirin, and penciclovir.

In parallel, the triphosphate forms of ribavirin (RBV-TP) and penciclovir (PEV-TP) were also able to inhibit the RdRp primer extension activity. Surprisingly, RBV-TP exerted the most potent inhibition on the poly-G template(Fig.1C), in contrast to the proposed model of ribavirin as a guanosine analog(*18*). PEV-TP showed comparable inhibition on poly-U and poly-C templates (Fig.1D), suggesting that guanine base can form mismatched pairing with pyrimidine base. The potency of PEV-TP is in the same range of FAV-TP and RBV-TP. To corroborate these biochemical data, we tested the ability of favipiravir, ribavirin, and penciclovir to inhibit viral replication in Vero E6 cells, in which the nucleotide prodrugs can be metabolized into the active triphosphate forms. In these cell-based assays, favipiravir, ribavirin, and penciclovir exhibited similar antiviral activity (Fig. 1E) with EC_50_ values of 96, 40, and 12 μM, respectively. In comparison, remdesivir has an EC_50_ value of 1.57 μM in cell-based anti-viral assays (Fig. 1E), suggesting remdesivir is more potent than favipiravir, ribavirin, and penciclovir. However, remdesivir has higher cell toxicity (CC_50_=264 μM) than favipiravir, ribavirin, and penciclovir, all three of them have CC_50_ values higher than 1000 μM (Fig. 1E).

To determine the structural basis of RdRp inhibition, we prepared the purified nsp12-7-8 complex (fig. S1A-1B) in the presence of an RNA template-primer duplex (fig. S1D) and the triphosphate forms of nucleotide inhibitors, which were added in more than 300 fold of molar excess to the RdRp to obtain a stable and completely inhibited complex for cryo-EM studies. For the FAV-bound RdRp complex, we collected over 3,209 micrograph movies of more than 5.303 million particle projections. Of these, 116,511 particles were used to yield a density map of 2.7 Å resolution (fig. S4). For the RBV-bound RdRp complex, we collected over 4,297 micrograph movies of more than 4.974 million particle projections. Of these, 196,511 particles were used to yield a density map of 2.97 Å resolution (fig. S5). For the PEV-bound RdRp complex, we collected over 3,913 micrograph movies of more than 5.240 million particle projections. Of these, 142,242 particles were used to yield a density map of 2.6 Å resolution for the overall conformation, and 139,773 particles were used to yield a density map of 2.74 Å resolution for the extended conformation (fig. S6). Because of the relatively high resolutions of all three structures, the key components of the RdRp complexes are well defined by clear EM density maps (Fig. 2A-AC, 2D-2E and fig. S7-9), including the RNA-template-primer duplex, one molecule of nsp12 and nsp7, and two molecules of nsp8.

**Figure 2.**
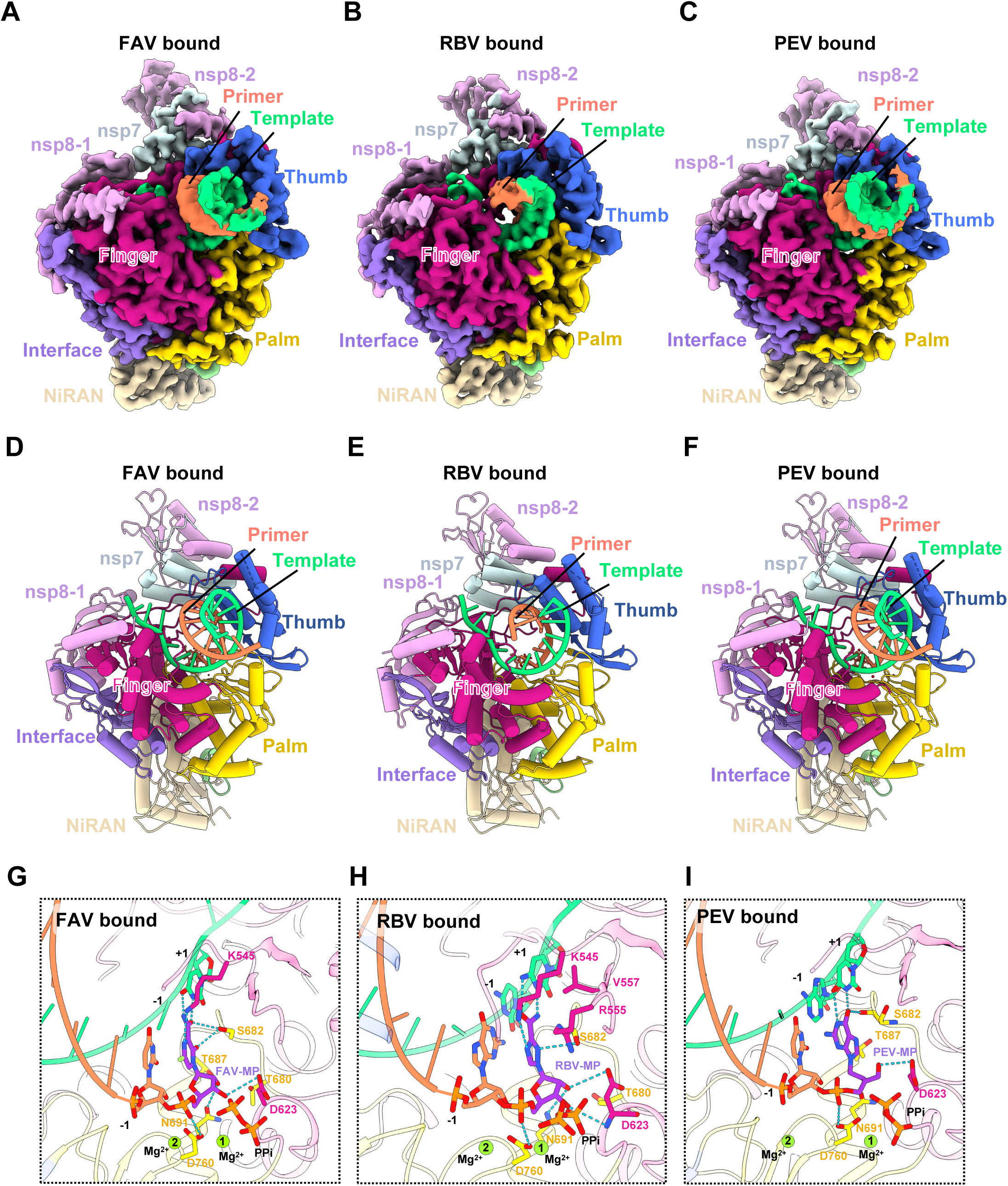
Cryo-EM Structures of the RdRp complex with RNA and nucleotide inhibitors. A-F, the cryo-EM map and structure of the nsp12-nsp7-nsp8-template-primer RNA complex bound to FAV (A and D), RBV (B and E), and PEV (C and F). G-I, A close view of the RdRp active site, showing the covalently bound FAV-MP (G), RBV-MP (H), PEV-MP (I), pyrophosphate, and magnesium ions.

The overall structures of the three nucleotide inhibitor-RdRp complexes are similar to each other with the RMSD values less than 0.3 Å for all Cα atoms of nsp12 (fig. S10). All three structures also resemble the previous remdesivir bound RdRp complex but the N-terminal NiRAN domain of nsp12 was completely resolved in the current three structures (fig. S7E, S8E and S9E). For the PEV-TP bound RdRp complex, the N-terminal segments of both nsp8 form two extended helices, acting like a pair of chopstick that clamp on the upstream RNA double strand (fig. S11), which is similar to the recent structures of RdRp in a transcribing conformation(*10, 19, 20*). For all three structures, the nucleotide inhibitors are in their monophosphate forms (FAV-MP, RBV-MP, and PEV-MP), which are located at the catalytic active site and covalently linked to the 3’ end of the primer strand (Fig. 2G-I). In addition, a pyrophosphate and two magnesium ions are observed at the active site in all three structures (Fig. 2G-I and fig. S7G, S8G and S9G), similar to the previous remdesivir-bound structure(*9*).

In addition to the covalent link of the monophosphate forms of the nucleotide inhibitors to the primer strand, the binding poses of these nucleotide inhibitors are stabilized by base pairing and stacking interactions with the double strand RNA, as well as interactions with protein residues at the catalytic center (Fig. 2G-I). For the FAV-bound RdRp complex, FAV-MP mimics an adenosine analog to form two hydrogen bonds with +1 uridine from the template strand (Fig. 2G). In addition, the FAV base forms hydrogen bond with K545 and S682 of nsp12 whereas the phosphor-ribose makes interactions with D623, N691 and the catalytic D760 (Fig. 2G). For the RBV-bound RdRp complex, RBV-MP mimics a guanosine analog to form two hydrogen bonds with +1 cytosine from the template strand (Fig. 2H). In addition, the RBV base forms hydrogen bond with K545 and S682 of nsp12, and stacking interaction with R555, whereas the phosphor-ribose makes interactions with D623, N691 and the catalytic D760, (Fig. 2H). For the PEV-bound RdRp complex, the guanosine from PEV-MP forms two hydrogen bonds with +1 mismatched uridine from the template strand (Fig. 2I). In addition, the guanosine base forms hydrogen bonds with S682 main chain CO of nsp12 whereas the phosphate backbone makes interactions with the catalytic D760 (Fig. 2I). The hydroxyl of penciclovir forms additional hydrogen bonds the side chain of D623 (Fig. 2I).

RNA replication by the viral RdRp involves at least two critical conformation states(*10, 21*), the pre-translocated conformation state in which the newly incorporated nucleotide is the +1 position within the catalytic site and the post-translocated conformation state in which the newly incorporated nucleotide is translocated into the -1 position, which will empty the +1 position to allow a new incoming nucleotide into the catalytic center. Currently the structures of SARS-CoV-2 RdRp have been solved with four different nucleotide inhibitors, FAV, RBV, PEV, and RDV. Comparison of these inhibitor-bound structures with the RdRp structure with native RNA(*10, 19, 20*) reveals that these nucleotide inhibitors are located at +1 position in the pre-translocated conformation state despite that FAV-TP, RBV-TP, and PEV-TP were added in vast excess during the complex formation (Fig. 3A). The positions of the base analogs, ribose and phosphate backbone of FAV-MP, RBV-MP, and RDV-MP are nearly overlapped with each other (Fig. 3A). PEV-MP does not have a ribose but its guanine base and phosphate backbone are also overlapped with FAV-MP and RBV-MP. The concentrations of the triphosphate form of these nucleotide inhibitors used in complex assembly were at 1 mM, which are comparable to the in vivo concentrations of natural NTP, thus it is surprised to observed that only a single base of nucleotide inhibitor was added to the 3’ end of the primer strand despite the availability of the 3’ OH from the nucleotide inhibitors to couple with the incoming 5’ triphosphate of the next inhibitor. Therefore, these inhibitors act like immediate chain terminators in the experimental conditions we used. Our structures are different from a recent structure of the RdRp in complex with RDV-TP in the presence of excess amount of GTP (Fig. 3B and fig. S12A), in which RDV-MP is at the -1 position and only one GMP at the +1 position despite the +2 position is a cytosine in the template strand, which is compatible with an incoming GTP. Thus, this inhibitor-RdRp complex is also stalled at the pre-translocated conformation despite an excess amount of native NTP for primer extension. This is in contrast to the RdRp RNA complex in the presence of only natural NTP (Fig. 3C-3E and fig. S12B-S12D), which structure reveals a post-translocated conformation, readily to adopted a new nucleotide for primer extension. Despite these structural differences, these nucleotide inhibitors seem to act through a common mechanism by stalling the RdRp complex at the stage of pre-translocated conformation.

**Figure 3.**
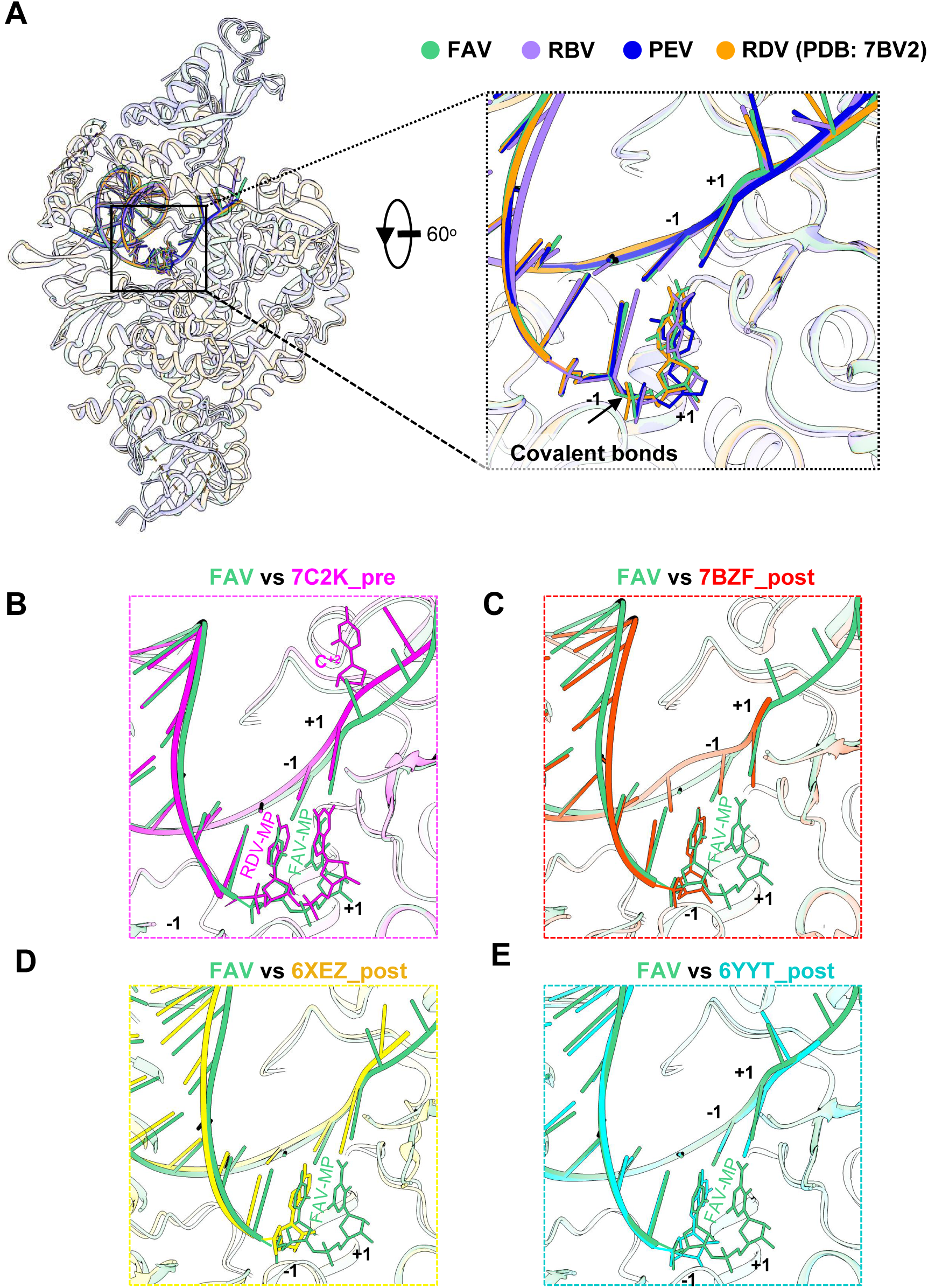
A common inhibition mechanism of nucleotide inhibitors. A, Comparisons of FAV/RBV/PEV bound SARS-CoV-2 RdRp complex with the RDV bound SARS-CoV-2 RdRp complex (PDB ID: 7BV2), with a close view of the RdRp active site, showing nucleotide inhibitors are covalently linked at the +1 site of the primer RNA strands. B-E close view of the RdRp active site from comparisons of FAV bound SARS-CoV-2 RdRp complex with the RDV bound SARS-CoV-2 RdRp complex (PDB ID: 7C2K, B), RNA bound SARS-CoV-2 RdRp complex (PDB ID: 7BZF, C), RNA bound SARS-CoV-2 RdRp complex (PDB ID: 6XEZ, D), RNA bound SARS-CoV-2 RdRp complex (PDB ID: 6YYT, E).

Among the above four nucleotide analog drugs, remdesivir is the first one that has been approved by FDA for COVID-19 through vein injection. This is probably that remdesivir has the most potent EC_50_ against SARS-CoV-2 in cell-based assays(*22*) (EC_50_ of remdesivir is ∼1-2 µM vs 10-100 µM for FAV, RBV, and PEV, Fig. 1E). However, remdesivir has disadvantages of difficulties in chemical synthesis and obligatory intravenous (IV) injection that requires hospital settings. In addition, pharmacokinetics of remdesivir shows that it is rapidly metabolized into its nucleotide form and it is highly enriched in liver that can cause hepatotoxicity(*23, 24*). Thus, it has been argued that parent nucleoside of remdesivir should have advantage for COVID-19 treatment(*25*). Indeed, cell-based inhibition assays of viral replication reveals that the parent nucleoside is about 3-fold more potent than remdesivir (Fig. 1E and Fig. 4A-B)(*22*). The deuterated analog bearing a deuterium atom at the C5 position of the base moiety further improve oral bioavailability (Fig. 4A and 4C). Prodrug modification of the deuterated nucleoside that is esterified with isobutyric acid further improved the viral inhibition potency (Fig. 4A and 4B). Together, the final modified compound X3 has an overall 5-fold potency improvement in inhibiting the viral replication over the original remdesivir (Fig. 4A and 4B).

**Figure 4.**
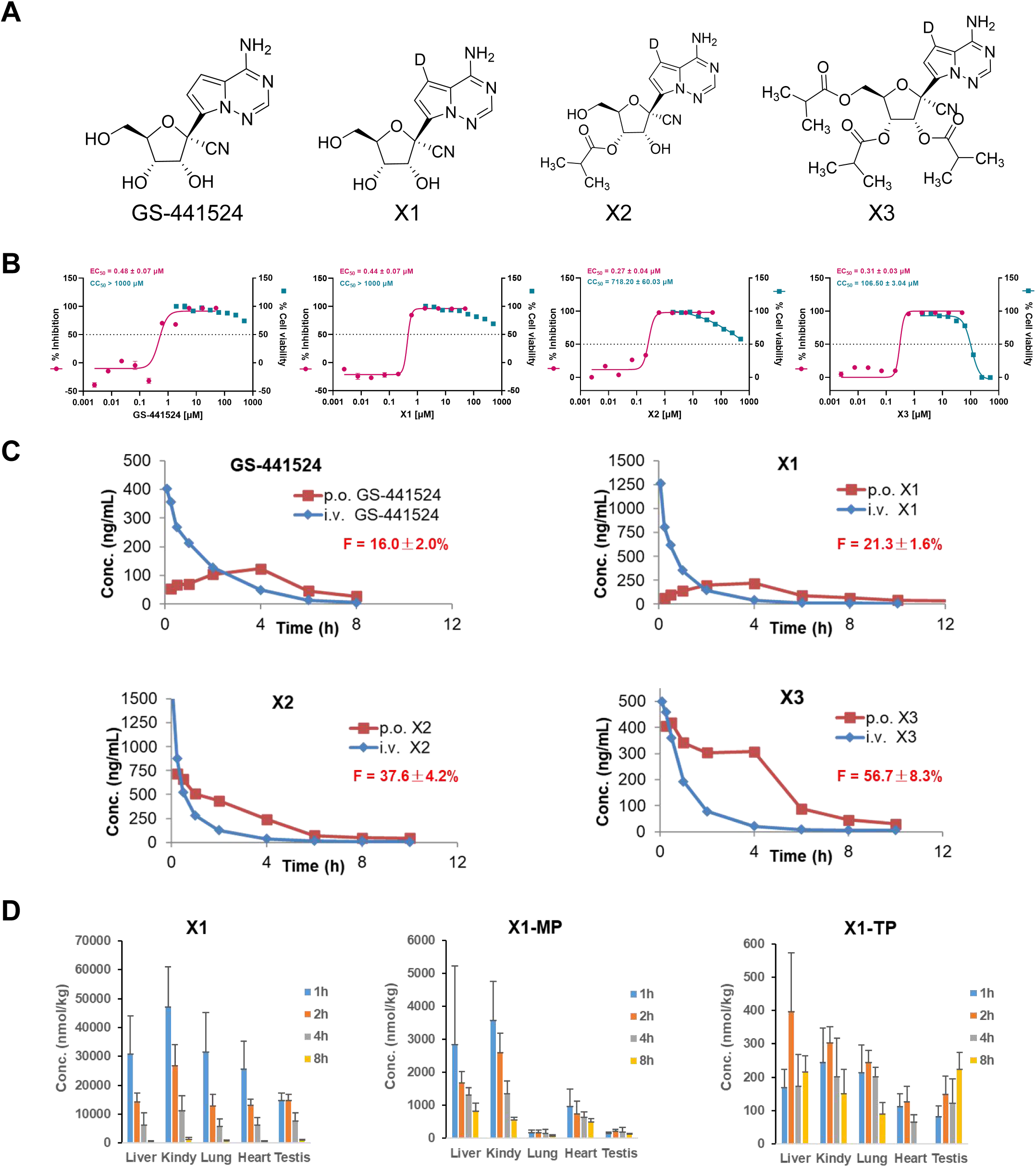
Modifications of GS-441524 provided novel and promising anti-SARS-CoV-2 nucleotide analogs. (A) The chemical structures of GS-441524, the deuterated derivative X1 and two isobutyrate prodrugs (X2 and X3) of X1. (B) Inhibition of SARS-CoV-2 replication and cellular toxicity of GS-441524, X1, X2 and X3. (C) Mean plasma concentrations of the corresponding nucleosides, following single intravenous (2 mg/Kg) and oral (10 mg/Kg) administration to SD rats (D)Tissue Distribution of X1, X1-MP, and X1-TP after oral administration of X3 to CD-1 mice at a single dose of 200 mg/kg (n = 5, Mean ± SD).

More importantly, compound X3 can be delivered via oral route with excellent bioavailability up to 56% (X3, Fig. 4C) and good retention time. In contrast to liver-enriched remdesivir, X3 achieves a well distribution of the nucleoside and nucleotide metabolite in the blood plasma as well as in many critical organs including lung that is vulnerable to SARS-CoV-2 infection, suggesting that X3 is a good drug candidate for treating COVID-19. A pilot experiment of inhibition of SARS-CoV-2 in mice transfected with human ACE2 demonstrated that mice treated with an acid salt form of X3 (development code: VV116) had 50-fold lower in viral load than the untreated mice (Fig S13A-C). The lung tissues of untreated mice were positive in immune staining of viral antigen of N protein but were negative in lung tissues from VV116-treated mice (Fig S13D). Together, these initial data demonstrated the antiviral efficacy of VV116 in the mouse model transfected with human ACE2.

The ongoing COVID-19 pandemic by no-stop infections of SARS-CoV-2 has caused a global health crisis that urgently request for an effective treatment. Because drug development normally takes a decade long of preclinical and clinical research, which is clearly unfit for current medical emergency. Drug repurposing that could bypass regular preclinical and clinical research is of critical importance in the current situation to combat the pandemics of COVID-19. In this paper, we demonstrated three existing drugs, favipiravir, ribavirin, and penciclovir, all nucleotide analogs, are inhibitors of the SARS-CoV-2 RdRp, an essential enzyme for the viral replication. We have further determined the structures of these three nucleotide analog drugs in complex with the SARS-CoC-2 RdRp and a template-primer RNA. Despite that penciclovir does not have a ribose scaffold that is shared by favipiravir and ribavirin, all nucleotide inhibitors are covalently linked to the primer strand in a monophosphate form, in which the RdRp RNA template-primer complex is stalled at the pre-translocated conformation state, revealing a common RdRp-inhibition mechanism that is also shared by the remdesivir bound RdRp complex. In addition, we discovered a terminal nucleotide transferase activity of the SARS-CoV-2 RdRp, which can add an additional nucleotide to the primer strand beyond the template end. The SARS-CoV-2 RdRp also display a remarkable promiscuity with respect to the incorporation of nucleotides that are unmatched with bases from the template strand. Based on these structures and previous remdesivir-bound structures, we designed several orally-available remdesivir derivatives that showed potent inhibition of viral replication in cell-based assays and animal models of SARS-CoV-2 infections. Our results provide a rational basis for repurpose of favipiravir, ribavirin, and penciclovir for treating COVID-19. The structures also provide a set of templates for designing the next generation of orally-available nucleotide analog drugs for COVID-19.

## Acknowledgments

The cryo-EM data were collected at the Cryo-Electron Microscopy Research Center, Shanghai Institute of Materia Medica. This work was partially supported by National Key R&D Program of China (2020YFC0861000), CAMS Innovation Fund for Medical Sciences No. 2020-I2M-CoV19-001, and Tsinghua University-Peking University Center for Life Sciences 045-160321001 to S. Z.; the National Key R&D Programs of China 2018YFA0507002; Shanghai Municipal Science and Technology Major Project 2019SHZDZX02 and XDB37030103 to H.E.X.; Zhejiang University special scientific research fund for COVID-19 prevention and control E33 and the National Science Foundation of China 81922071 to Y.Z.; the 2020 ANSO Collaborative Research Project (ANSO-CR-SP-2020-03) to J. S.; the 100 Talents Program of the Chinese Academy of Sciences, Chinese Academy of Sciences grant (XDA12010317), Natural Science Foundation of Shanghai (18ZR1447700) to X.Y; the National Natural Science Foundation of China (31900869) and Shanghai Sailing Program (19YF1456800) to Z.L; National Science & Technology Major Project “Key New Drug Creation and Manufacturing Program” of China (2018ZX09711002), Science and Technology Commission of Shanghai Municipal 20431900100 and Jack Ma Foundation 2020-CMKYGG-05 to H.J.; National Natural Science Foundation 31770796, National Science and Technology Major Project 2018ZX09711002, and K.C. Wong Education Foundation to Y.J, the National Natural Science Foundation of China (31970165) to L. Z.

## Author contributions

W.Y. designed the expression constructs, purified the RdRp complex, prepared samples and cryo-EM grids, performed data collection and processing toward the structure determination, analyzed the structures, and prepared figures and manuscript. X.L. designed RdRp activity assays, participated in data interpretation and figure preparation; Z.L., Z.Z., Q. W., and X.Y. evaluated the specimen by negative-stain EM, screened the cryo-EM conditions, prepared the cryo-EM grids and collected cryo-EM images, performed data processing, density map calculations, model building, and figure preparation; Y.Z. and D.-D.S, analyzed EM data; Y.X., G.T., H.A.A., T.H., D.W., and J.S. designed and prepared nucleotide derivatives; J.L. performed PK of nucleotide derivatives; L.Z., Q.W., and G.X. designed and performed cell-based viral inhibition assays; X. W., F. Z., and M.G. participated in expression, purification and functional assays of the RdRp; Y.J. participated in experimental design and manuscript editing; H.J. conceived and coordinated the project; S.Z. conceived the project, initiated collaboration with H.E.X. and supervised X.L.; H.E.X. conceived and supervised the project, analyzed the structures, and wrote the manuscript with inputs from all authors.

## Data and materials availability

Density maps and structure coordinates have been deposited with immediate release. The accession numbers of Electron Microscopy Database and the Protein Data Bank are EMD-XXXX and PDB ID XXXX for the FAV-RdRp complex; EMD-XXX and PDB ID XXXX for the RBV-RdRp complex; EMD-XXX/ and PDB ID XXXX/ for the two PEV-RdRp complexes. Materials are available up on request.

## Supplementary figures

**Figure S1.**
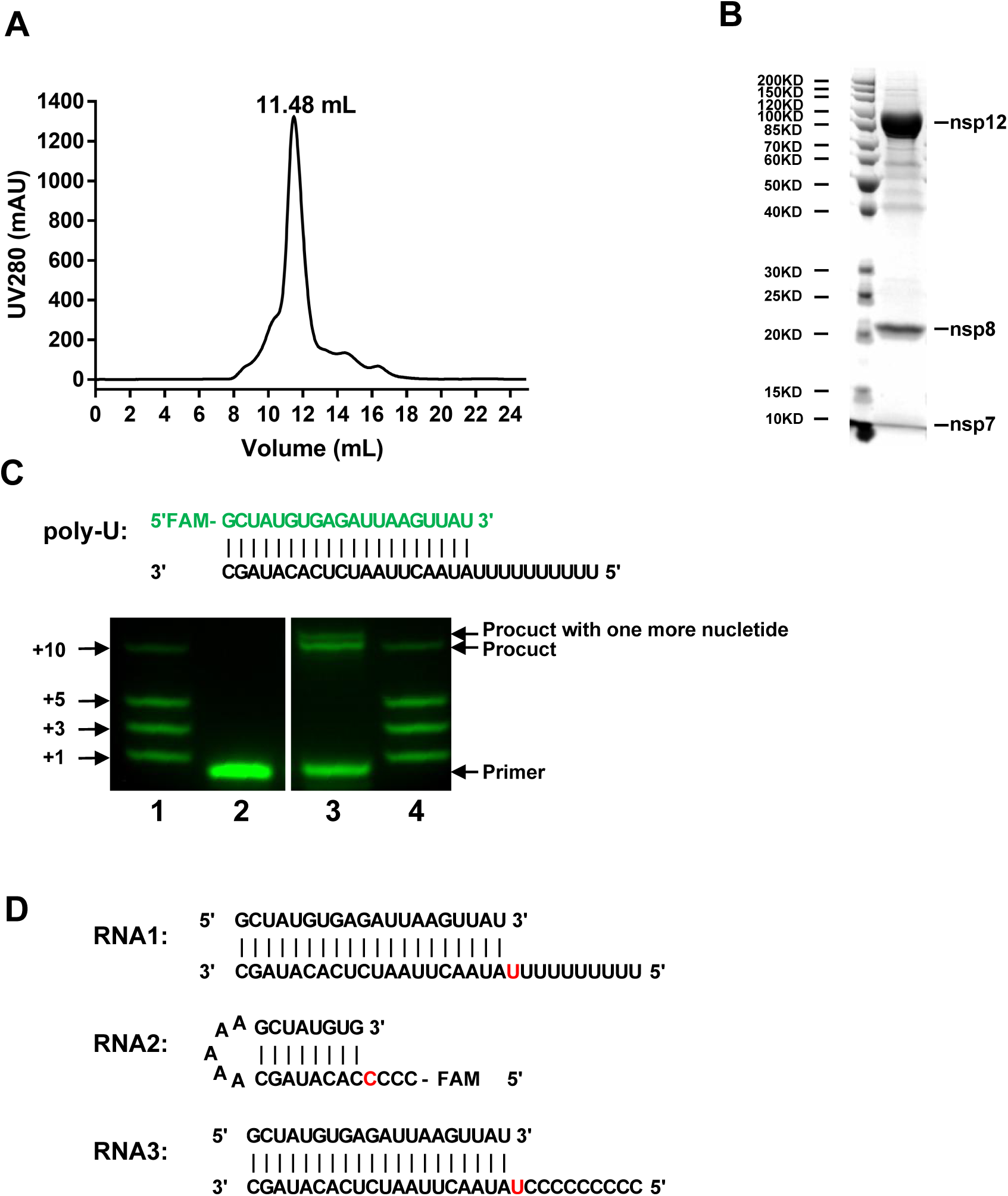
Purification and characterization of the RdRp complex. A, Gel filtration profile of the RdRp complex with additional nsp7 and nsp8, showing a sharp peak. B, SDS gel of the purified RdRp complex with additional nsp7 and nsp8, showing balanced ratios for each subunit. C, Gel-based elongation assay with the 20% Urea-PAGE denatured gel to observe a product with one nucleotide longer than the template. Lane 1 and Lane 4 shows a mixture of synthesized RNA strands with the indicated lengths of adenosine nucleotides added into the primer strand of 5-FAM-GCUAUCUCAGAUUAAGUUAU as the marker, lane 2 indicates the primer strand of 5-FAM-GCUAUCUCAGAUUAAGUUAU as control, lane 3 indicates the product with one nucleotide longer than the template. D, Diagram of the template and primer duplex RNAs used in the RdRp structural studies. RNA1 was used in the FAV-bound RdRp complex; RNA2 for the RBV-bound RdRp complex; and RNA3 for the PEV-bound RdRp complex.

**Figure S2.**
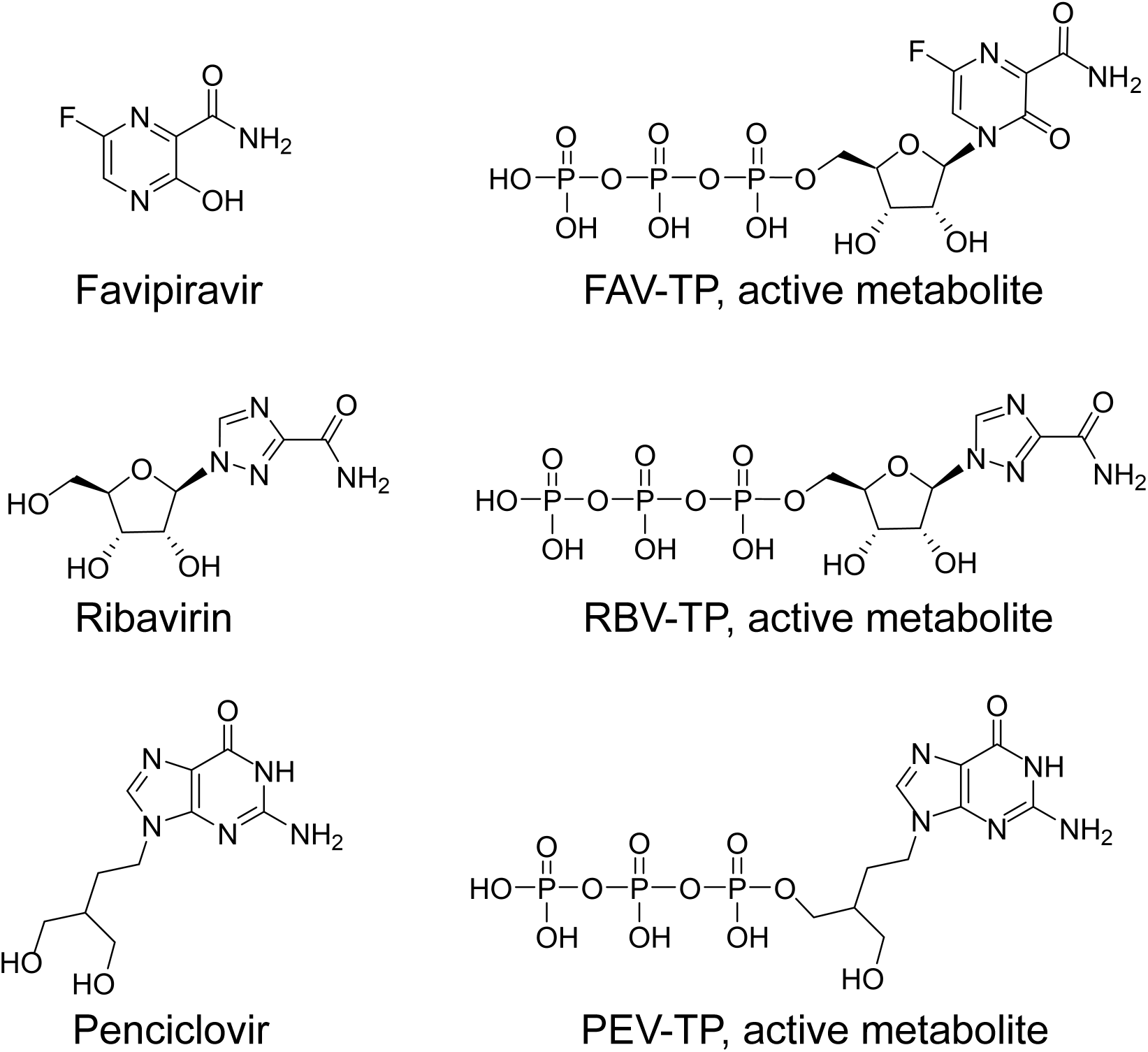
The chemical structures of prodrug forms and triphosphate forms of favipiravir, ribavirin, and penciclovir.

**Figure S3.**
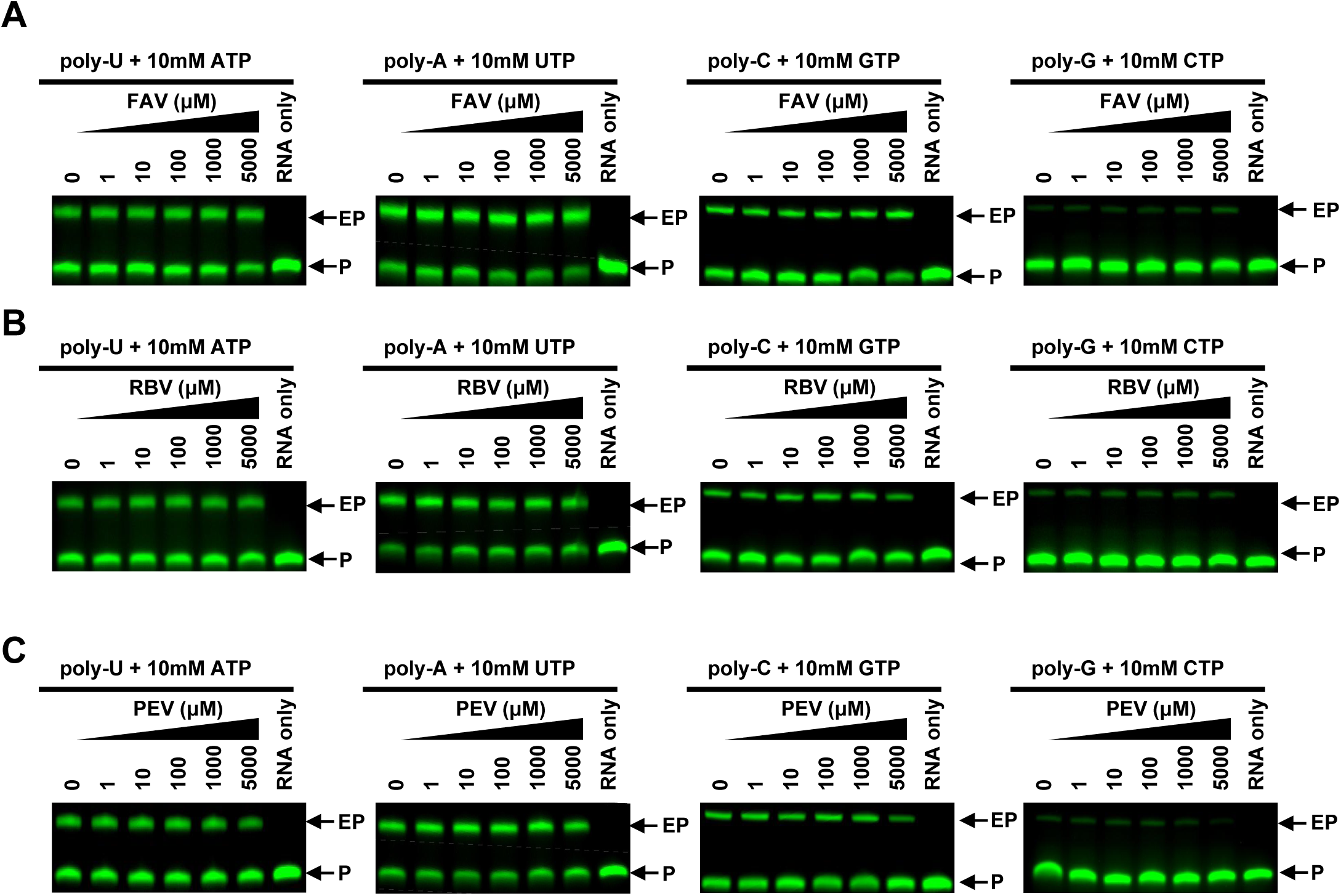
No inhibitions of the different RNA substrates by the prodrug forms of FAV (A), RBV (B), and PEV (C) based on the gel-based elongation assays. The EP means the elongated product, while the P means the primer RNA strand.

**Figure S4.**
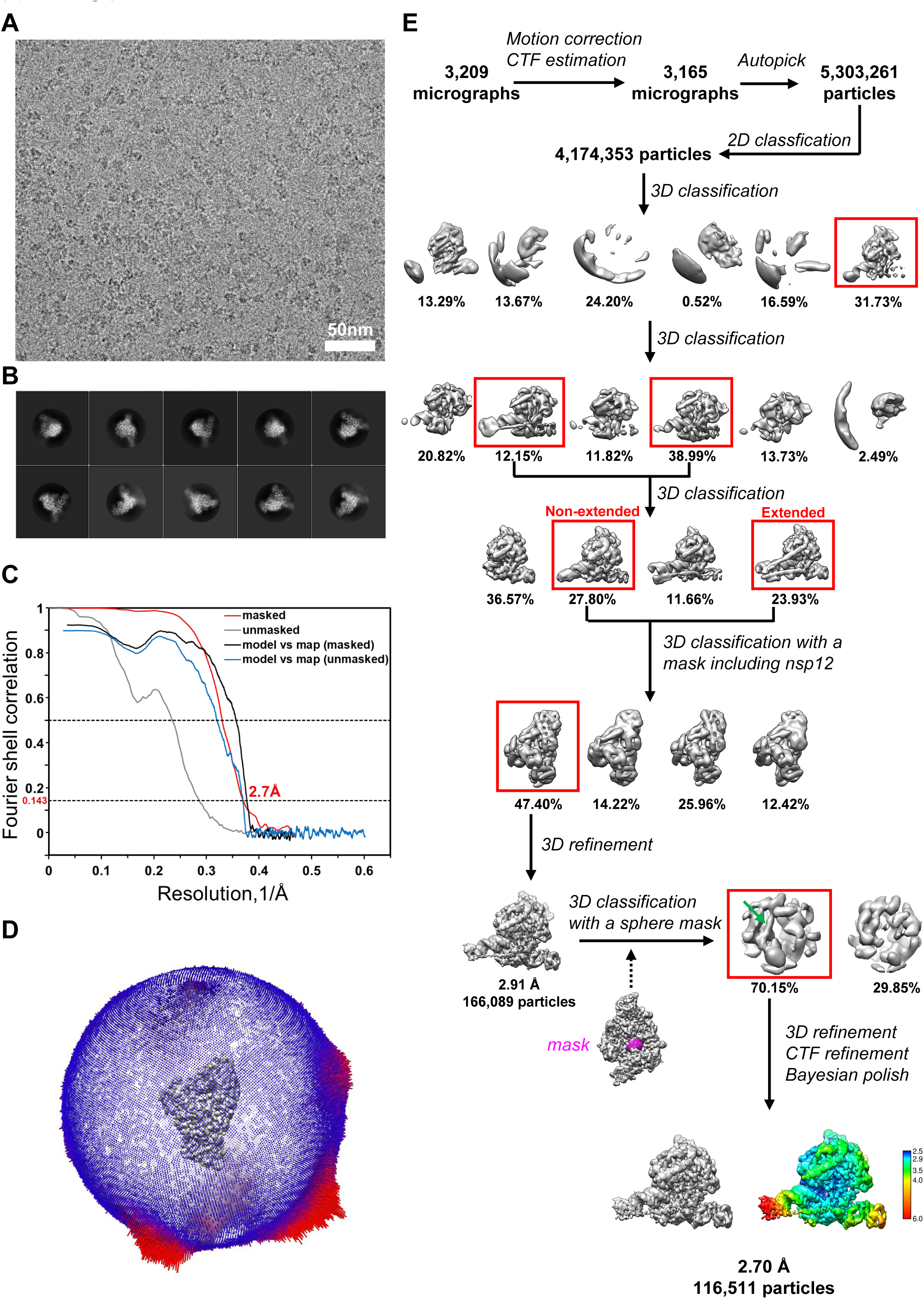
Single particle cryo-EM analysis of the FAV-bound RdRp complex. A, Representative cryo-EM micrograph of the FAV-bound RdRp complex. B, Representative 2D class averages of the FAV-bound RdRp complex. C, Fourier shell correlation curves of cryo-EM map for the FAV-bound RdRp complex. D, Euler angle distribution of particles used in the final reconstruction. E, Flowchart of cryo-EM works of the FAV-bound RdRp complex with maps colored by local resolution (Å).

**Figure S5.**
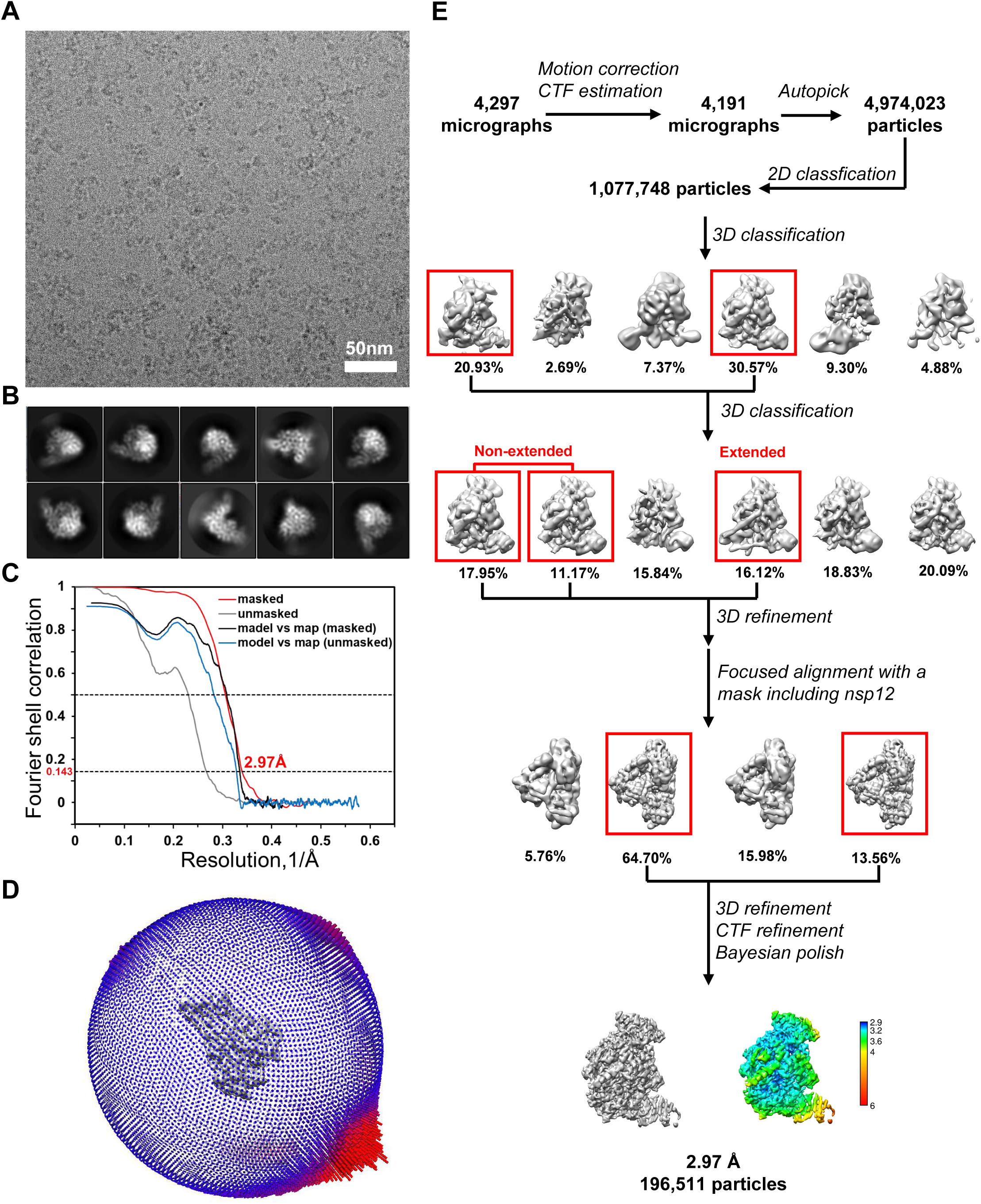
Single particle cryo-EM analysis of the RBV-bound RdRp complex. A, Representative cryo-EM micrograph of the RBV-bound RdRp complex. B, Representative 2D class averages of the RBV-bound RdRp complex. C, Fourier shell correlation curves of cryo-EM map for the RBV-bound RdRp complex. D, Euler angle distribution of particles used in the final reconstruction. E, Flowchart of cryo-EM works of the RBV-bound RdRp complex with maps colored by local resolution (Å).

**Figure S6.**
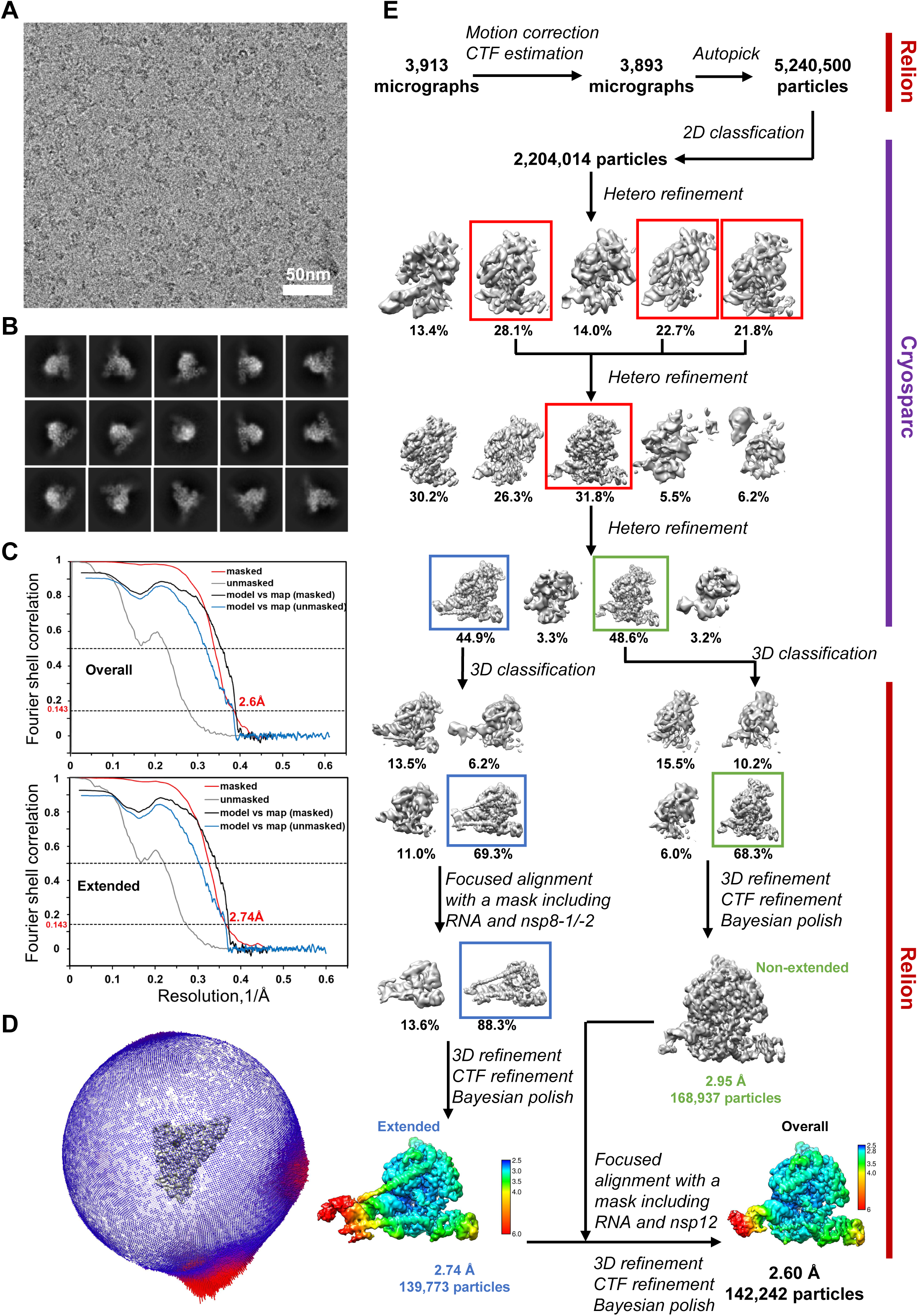
Single particle cryo-EM analysis of the PEV-bound RdRp complex. A, Representative cryo-EM micrograph of the PEV-bound RdRp complex. B, Representative 2D class averages of the PEV-bound RdRp complex. C, Fourier shell correlation curves of cryo-EM map for the PEV-bound RdRp complex. D, Euler angle distribution of particles used in the final reconstruction. E, Flowchart of cryo-EM works of the PEV-bound RdRp complex with maps colored by local resolution (Å).

**Figure S7.**
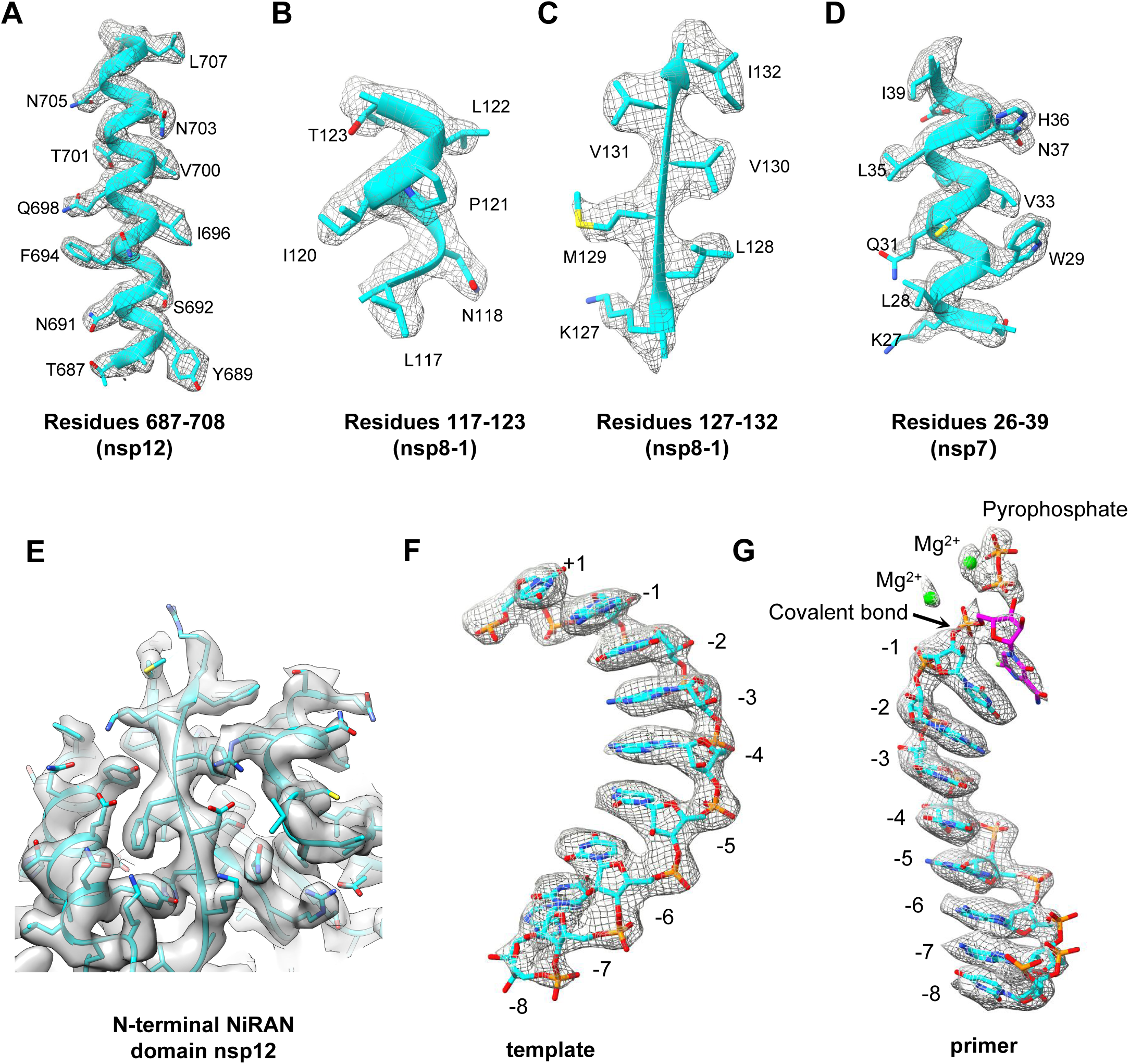
Cryo-EM map for the FAV-bound RdRp complex. A-G. Cryo-EM map and model of nsp12 (residues 687-708) (A), nsp8-1 (residues 117-123) (B), nsp8-1 (residues 127-132) (C), nsp7 (residues 26-39) (D), the N-terminal NiRAN domain of nsp12(E), the template strand RNA (position -8 to +1) (F), and the primer strand RNA, pyrophosphate, two magnesium ions and the covalently bound FAV-MP (G).

**Figure S8.**
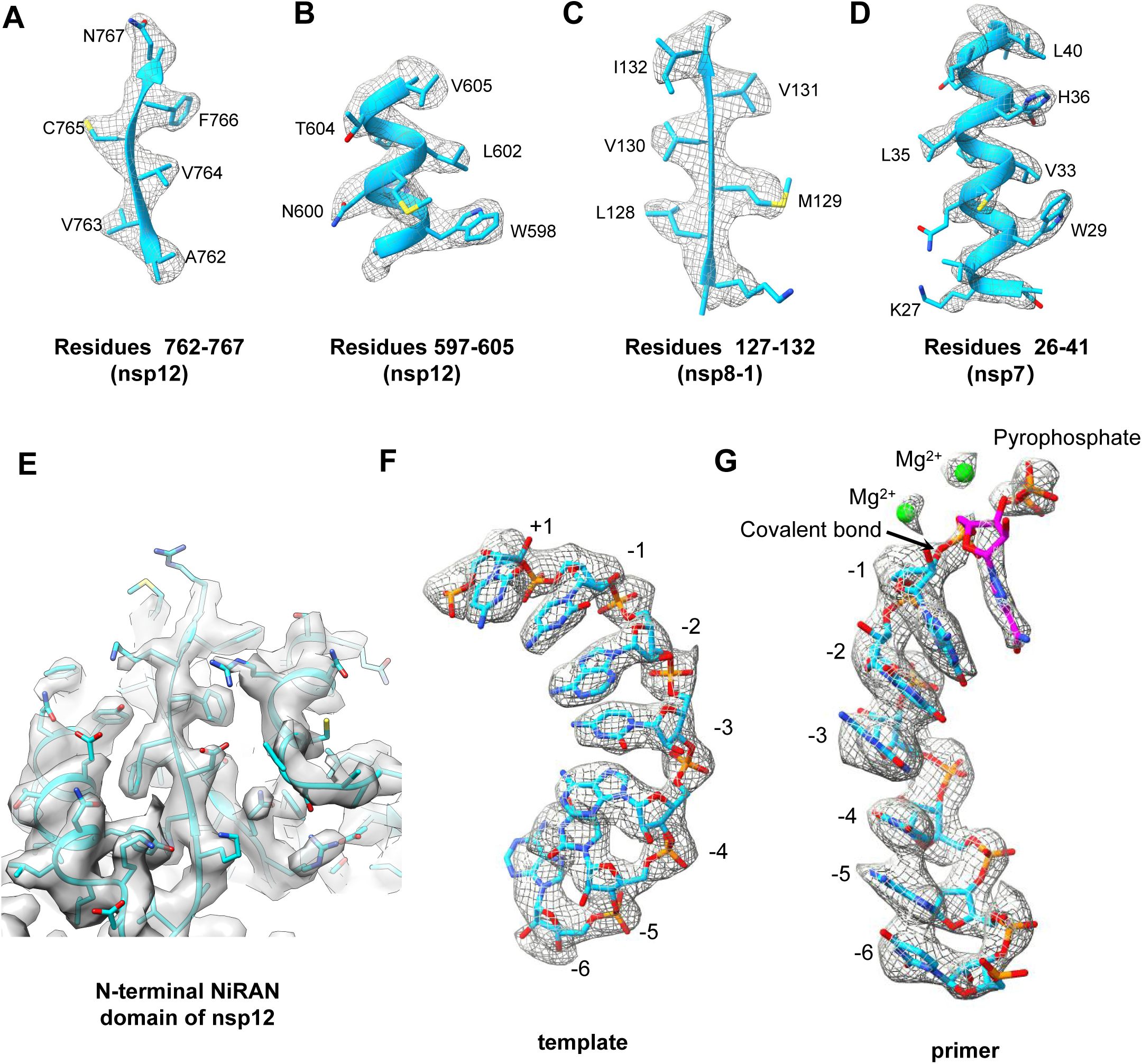
Cryo-EM map for the RBV-bound RdRp complex. A-G. Cryo-EM map and model of nsp12 (residues 762-767) (A), nsp12 (residues 597-605) (B), nsp8-1 (residues 127-132) (C), nsp7 (residues 26-41) (D), the N-terminal NiRAN domain of nsp12(E), the template strand RNA (position -6 to +1) (F), and primer strand RNA, pyrophosphate, two magnesium ions and the covalently bound RBV-MP (G.

**Figure S9.**
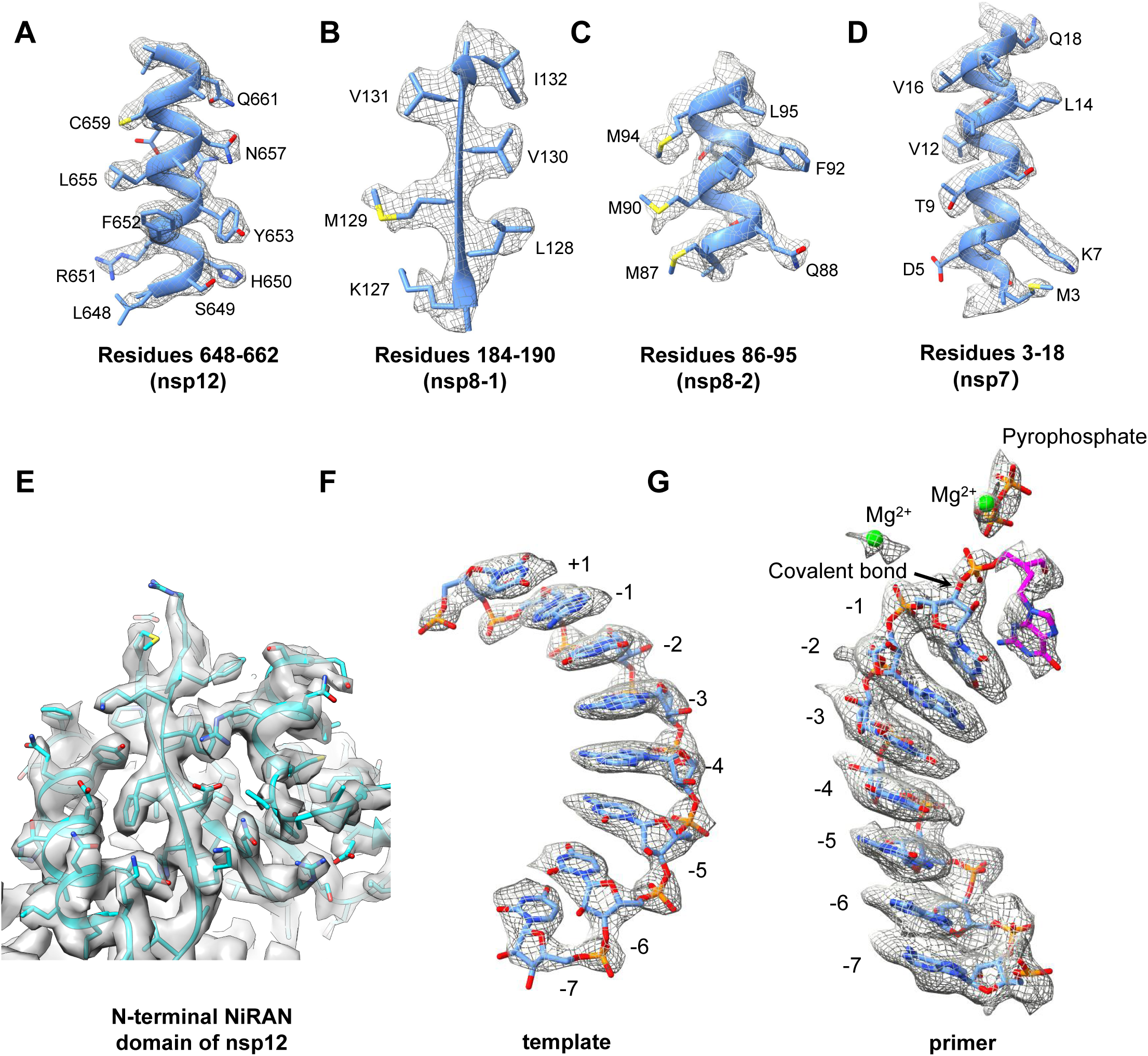
Cryo-EM map for the PEV-bound RdRp complex. A-G. Cryo-EM map and model of nsp12 (residues 648-662) (A), nsp8-1 (residues 184-190) (B), nsp8-2 (residues 86-95) (C), nsp7 (residues 3-18) (D), the N-terminal NiRAN domain of nsp12(E), the template strand RNA (position -7 to +1) (F), the primer strand RNA, pyrophosphate, two magnesium ions and the covalently bound PEV-MP (G).

**Figure S10.**
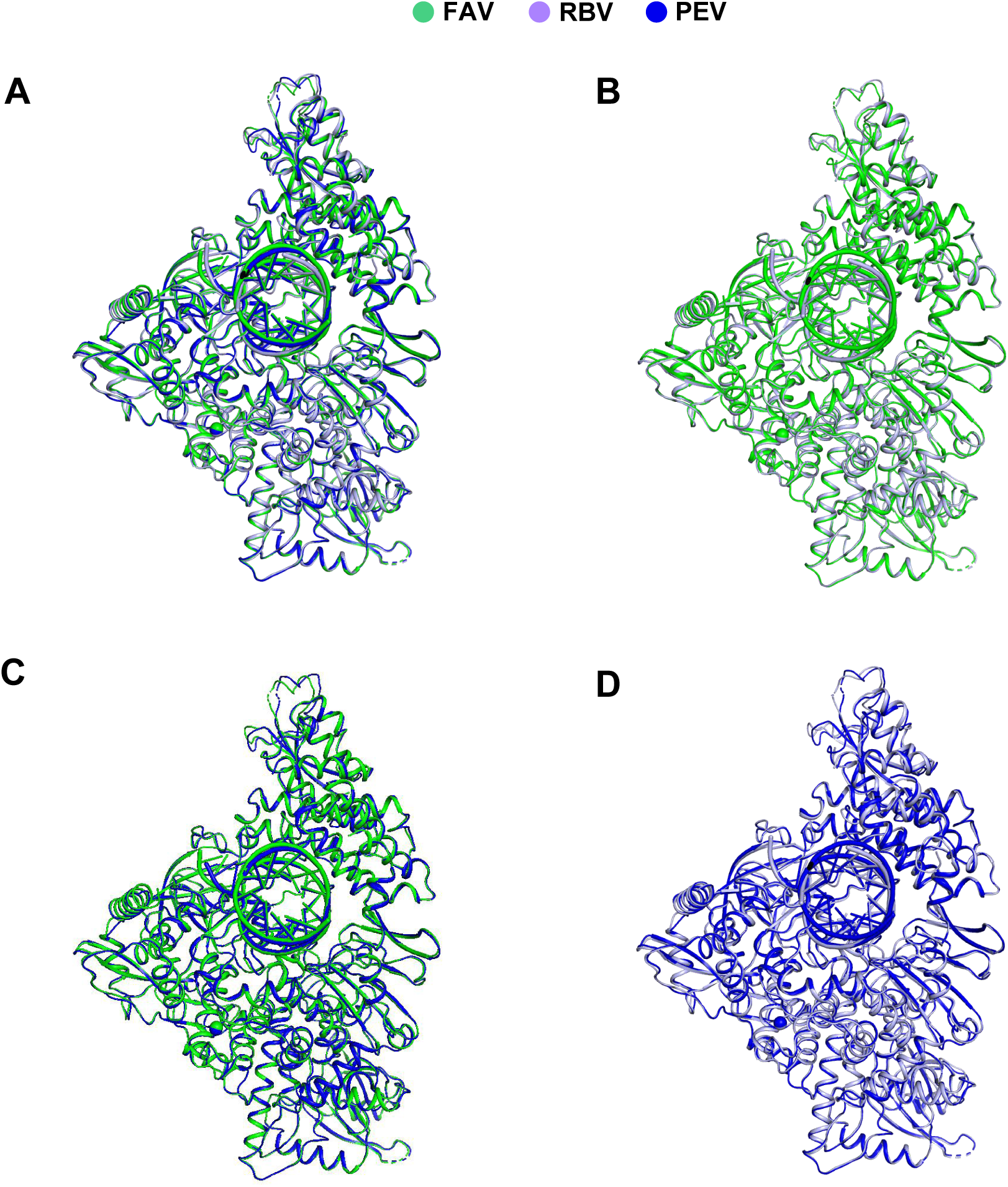
Comparisons of FAV/RBV/PEV bound SARS-CoV-2 RdRp complex. A, Comparison of FAV/RBV/PEV bound SARS-CoV-2 RdRp complex. B, Comparison of FAV/RBV bound SARS-CoV-2 RdRp complex. C, Comparison of FAV/PEV bound SARS-CoV-2 RdRp complex. D, Comparison of RBV/PEV bound SARS-CoV-2 RdRp complex.

**Figure S11.**
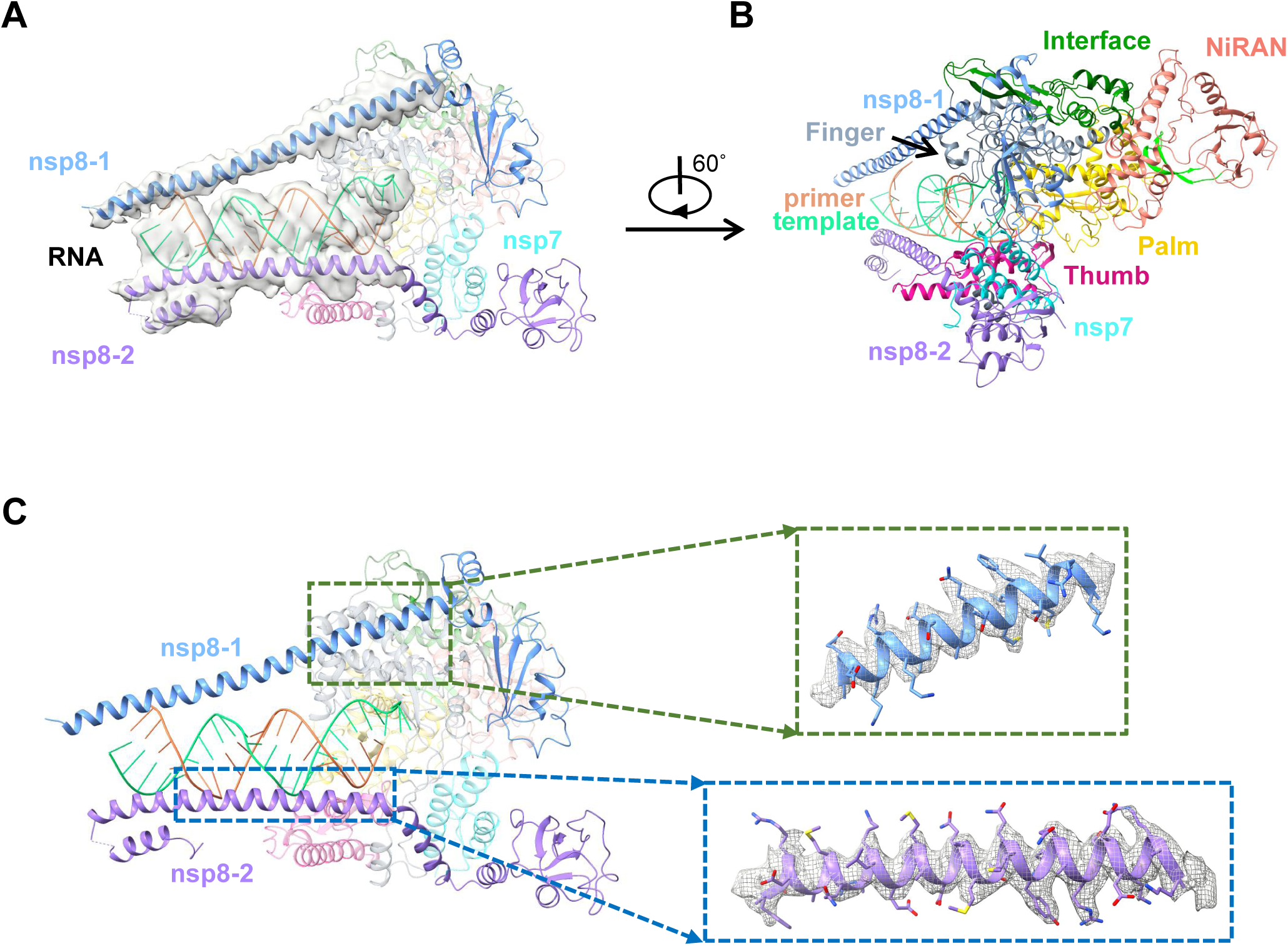
Cryo-EM Structure of the PEV-bound RdRp complex in nsp8-extended conformation. A-B. Two views of structure of PEV-bound RdRp complex in extended conformation. The cryo-EM density map for extended helices from two nsp8 copies and the clamped RNA duplex were shown in (A). C. Cryo-EM map and model of nsp8-1(residues 76-97) and nsp8-2(residues 49-82).

**Figure S12.**
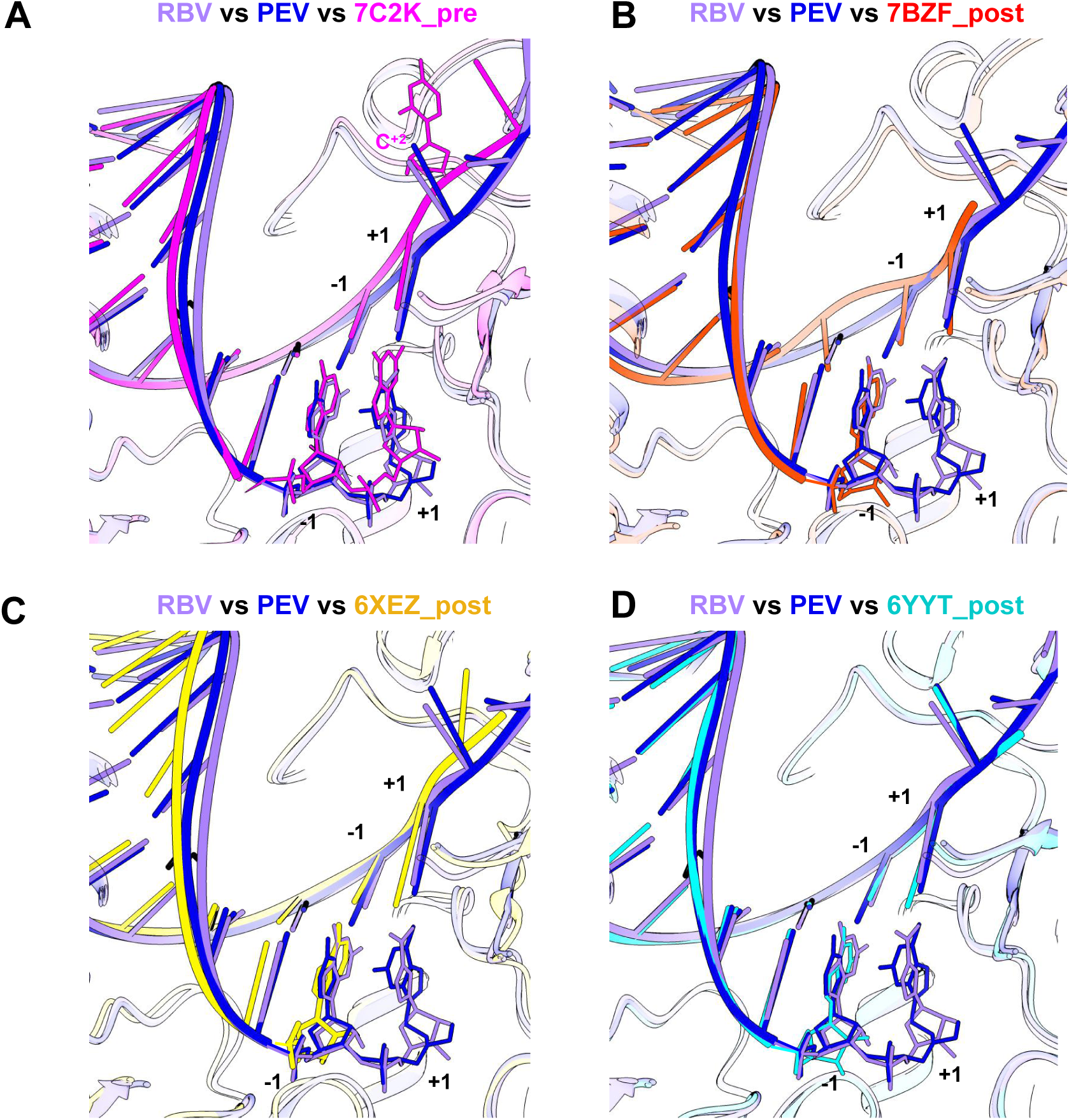
The similar conformations of RBV/PEV bound SARS-CoV-2 RdRp. A close view of the RdRp active site from comparisons of RBV/PEV bound SARS-CoV-2 RdRp complex with the RDV bound SARS-CoV-2 RdRp complex (PDB ID: 7C2K, A), RNA bound SARS-CoV-2 RdRp complex (PDB ID: 7BZF, B), RNA bound SARS-CoV-2 RdRp complex (PDB ID: 6XEZ, C), RNA bound SARS-CoV-2 RdRp complex (PDB ID: 6YYT, D).

**Figure S13.**
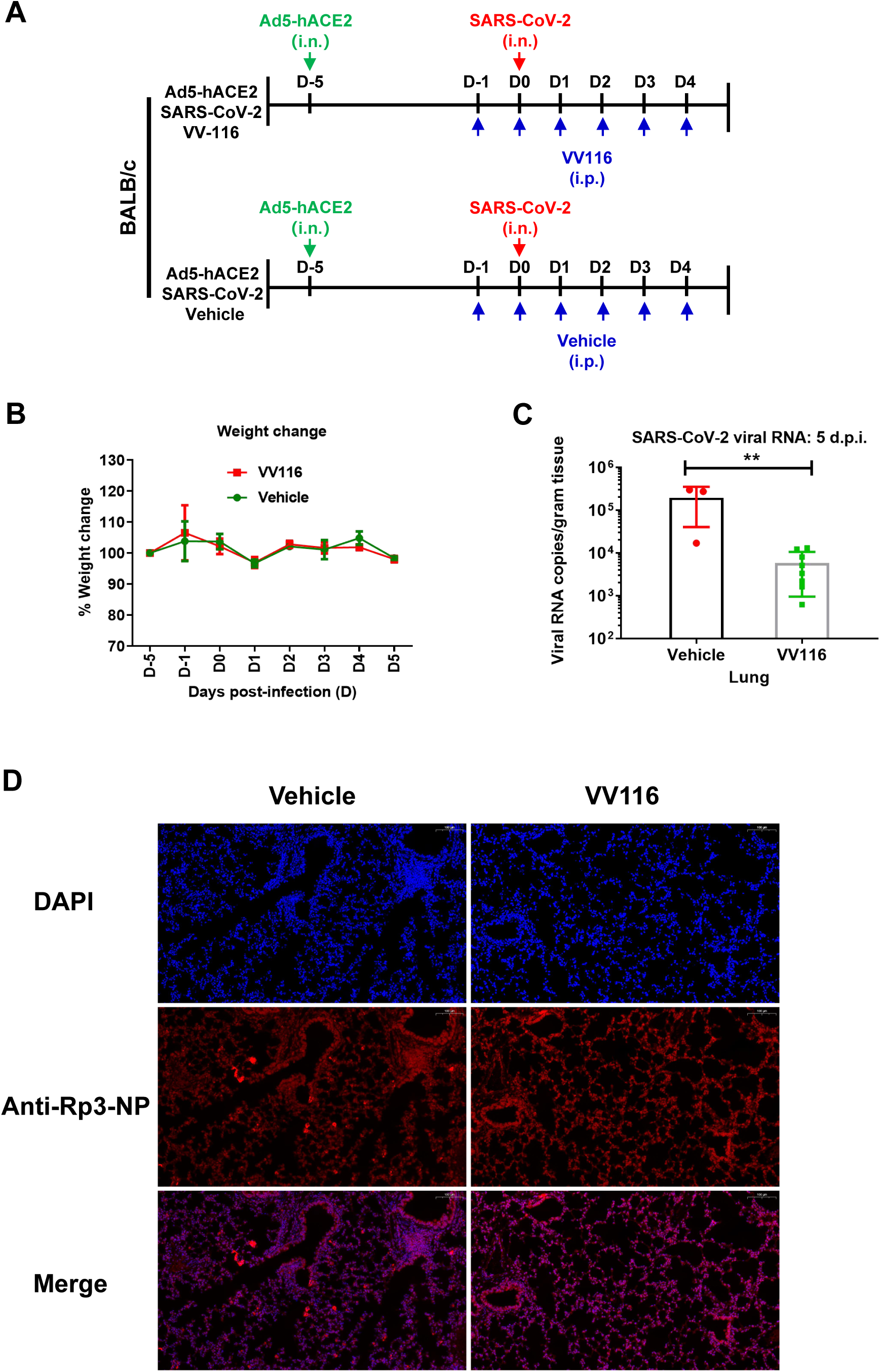
In vivo anti-SARS-CoV-2 efficacy of VV116 in mouse Ad5-hACE2 model. A. 6-to-8-week-old female BALB/c mice were intranasally infected with 50 μl 5×109 PFU/ml of Ad5-hACE2 (day -5), and were treated with VV116 (100 mg/kg, n=8) or vehicle (n=3). At day 0, mice were infected with 100,000 PFU of SARS-CoV-2 and treated with VV116 (100 mg/kg) or vehicle daily. Mice were sacrificed at day 5. B. Weight change was monitored. Error bars indicate SD. C. Viral RNA levels in lung tissues of vehicle-controlled and VV116-treated mice. Error bars indicate SD. D. Viral antigen was detected by anti-Rp3-CoV N protein polyclonal antibody (red) in lung tissue collected at day 5. Images were collected using FV1200 confocal microscopy. Scale bar is 100 μm.

**Table S1, EM Data and Structure Refinement Statistics** (shown in the last page after supplementary figures)

## Supplementary Materials

### Materials and Methods

#### Constructs and expression of the RdRp complex

The RdRp complex was prepared according to the same method reported(*17*) as described below. The full-length gene of the SARS-CoV-2 nsp12(residues 1-932) was chemically synthesized with codon optimization (General Biosystems). The gene was cloned into a modified pFastBac baculovirus expression vector containing a 5’ ATG starting sequence and C-terminal Tobacco Etch Virus (TEV) protease site followed by a His8 tag. The plasmid contains an additional methionine at the N-terminus and GGSENLYFQGHHHHHHHH at the C-terminus of nsp12. The full-length genes for nsp7 (residues 1-83) and nsp8 (residues 1-198) were cloned into the pFastBac vector containing a 5’ ATG starting sequence. All constructs were generated using the Phanta Max Super-Fidelity DNA Polymerase (Vazyme Biotech Co.,Ltd) and verified by DNA sequencing. All constructs were expressed in *Spodoptera frugiperda* (*Sf*9) cells. Cell cultures were grown in ESF 921 serum-free medium (Expression Systems) to a density of 2-3 million cells per ml and then infected with three separate baculoviruses at a ratio of 1:2:2 for nsp12, nsp7 and nsp8 at a multiplicity of infection (m.o.i.) of about 5. The cells were collected 48 h after infection at 27 °C and cell pellets were stored at − 80 °C until use.

In addition, the genes of nsp7 and nsp8 were cloned into a modified pET-32a(+) vector containing a 5’ ATG starting sequence and C-terminal His8 tag with a TEV cleavage site for expression in E. coli. Plasmids were transformed into BL21(DE3) (Invitrogen). Bacterial cultures were grown to an OD600 of 0.6 at 37 °C, and then the expression was induced with a final concentration of 0.1 mM of isopropyl β-D-1-thiogalactopyranoside (IPTG) and the growth temperature was reduced to 16 °C for 18-20 h. The bacterial cultures were pelleted and stored at − 80 °C until use.

#### Purification of the RdRp complex

The purification of nsp7 and nsp8 expressed in bacteria was similar to the purification of nsp7 and nsp8 reported previously(*17*). Briefly, bacterial cells were lysed with a high-pressure homogenizer operating at 800 bar. Lysates were cleared by centrifugation at 25,000 × g for 30 min and were then bound to Ni-NTA beads (GE Healthcare). After wash with buffer containing 50 mM imidazole, the protein was eluted with buffer containing 300 mM imidazole. The tag was removed with incubation of TEV protease overnight and protein samples were concentrated with 3 kDa or 30 kDa molecular weight cut-off centrifuge filter units (Millipore Corporation) and then size-separated by a SuperdexS75 10/300 GL column in 25 mM HEPES pH 7.4, 200 mM sodium chloride, 5% (v/v) glycerol. The fractions for the nsp7 or nsp8 were collected, concentrated to about 10 mg/ml, and stored at − 80 °C until use.

The insect cells containing the co-expressed RdRp complex were resuspended in binding buffer of 25 mM HEPES pH 7.4, 300 mM sodium chloride, 25 mM imidazole, 1 mM magnesium chloride, 0.1% (v/v) IGEPALCA-630 (Anatrace), 1 mM tris (2-carboxyethyl) phosphine (TCEP), 10% (v/v) glycerol with additional EDTA-free Protease Inhibitor Cocktail (Bimake), and then incubated with agitation for 20 min at 4 °C. The incubated cells were lysed with a high-pressure homogenizer operating at 500 bar. The supernatant was isolated by centrifugation at 30,000×g for 30 min, followed by incubation with Ni-NTA beads (GE Healthcare) for 2 h at 4 °C. After binding, the beads were washed with 20 column volumes of wash buffer of 25 mM HEPES, pH 7.4, 300 mM sodium chloride, 25 mM imidazole, 1 mM magnesium chloride, 1 mM TCEP, and 10% (v/v) glycerol. The protein was eluted with 3-4 column volumes of elution buffer of 25 mM HEPES pH 7.4, 300 mM sodium chloride, 300 mM imidazole, 1 mM magnesium chloride, 1 mM TCEP, and 10% (v/v) glycerol.

The co-expressed RdRp complex was incubated with additional nsp7 and nsp8 from the bacterial expression in a 1:1:2 molar ratios and incubated at 4 °C for 4h. Incubated RdRp complex was concentrated with a 100 kDa molecular weight cut-off centrifugal filter unit (Millipore Corporation) and then size-separated by a SuperdexS200 10/300 GL column in 25 mM HEPES pH 7.4, 300 mM sodium chloride, 0.1 mM magnesium chloride, 1 mM TCEP. The fractions for the monomeric complex were collected and concentrated. For the nucleotide inhibitor bound complex, the concentrated RdRp complex were diluted to 0.5 mg/ml with buffer of 25 mM HEPES pH 7.4, 100 mM sodium chloride, 2 mM magnesium chloride, 1 mM TCEP, combined with the corresponding template-primer RNA (fig. S1D) in a 1: 1.5 molar ratio and incubated at 4 °C for 0.5h. Then the triphosphate form of nucleotide inhibitor (FAV-TP, RBV-TP and PEV-TP as shown in fig. S2) was added to the RNA incubated RdRp complex at 1 mM concentration respectively, and incubated at 4 °C for another 0.5h. Then the incubated nucleotide inhibitor bound RdRp complex was concentrated up to 5-8 mg/ml for EM experiments.

#### Cryo-EM sample preparation and data acquisition

Aliquots of 3 μL samples of protein-RNA complexes at a concentration of 5-8 mg/mL containing 0.0035% DDM were applied onto glow-discharged 200-mesh grids (Quantifoil R1.2/1.3). Grids were blotted with filter paper for 3.0 s with a blot force of 1 and plunge-frozen in liquid ethane using a FEI Vitrobot Mark IV. Cryo-EM movies were collected on a 300kV Titan Krios microscope (FEI) equipped with a Gatan image filter (operated with a slit width of 20 eV) (GIF) and K3 direct detection camera. The microscope was operated at a calibrated magnification of 46,773 X, yielding a pixel size of 1.069 Å on micrographs. All movies were automatically collected by SerialEM(*26*) and the exposure time was set to 3s with an accumulated dose of ∼69 e^−^/Å^2^ on sample. For the sample of FAV-TP complex, 3,209 movies were collected with a defocus range of -0.4 μm to -2.5 μm. For the sample of PEV-TP complex, 3,913 movies were collected with a defocus range of -0.4 μm to -2.3 μm. For the sample of RBV-TP complex, 4,297 movies were collected with a defocus range of -0.7 μm to -2.6 μm. Each micrograph was fractionated into a movie stack of 36 image frames.

#### Image processing

Frames in each movie stack were aligned for beam-induced motion correction using the program MotionCorr2(*27*), and the resulting dose-weighted micrographs were used for the further process. CTFFIND4(*28*) was used to determine the contrast transfer function (CTF) parameters.

For the sample of PEV-TP complex, 3,893 good micrographs were selected after discarding the ones with crystal ice contamination. Auto-picking program of Relion 3.0(*29*) was used to pick the particles with the model of the Remdesivir-bound RdRp complex of SARS-CoV-2 (PDB ID: 7BV2)(*9*) as a reference, yielding a total of 5,240,500 picked particles. Then, the extracted particle stack was transferred to software Cryosparc v2(*30*) and a round of reference-free 2D classification was performed. 2,204,014 particles were selected from classes representing projections of protein-RNA RdRp complex in different orientations, and subjected to three rounds of Heterogenous Refinement using a reconstruction of the RDV-TP complex of SARS-CoV-2 (EMD-30210)(*9*) as a 3D template. In the third round of Heterogenous Refinement, one of the two high-resolution 3D averages displayed two prominent long helical extensions in the two copies of nsp8 (extended form) that clamp the upstream RNA duplex, which, however, are absent in the other one (non-extended form). Next, the particle sets of extended form and non-extended form are separately imported from Cryosparc v2 to Relion 3.0 and subjected to one another round of 3D classification. For the structure determination of PEV-TP complex in extended form, a round of focused alignment with a mask including RNA and nsp8-1/-2 was performed before 3D refinement, sorting out 139,773 particles with ordered helical extensions of nsp8-1/-2. After consecutive rounds of CTF refinement and Bayesian polishing of particles, another round 3D refinement was performed, yielding a final reconstruction at a global resolution of 2.74 Å. For the structure determination of PEV-TP complex in non-extended form, 168,937 selected particles were subjected to rounds of 3D refinement, CTF refinement, Baysian particle polishing, and finally generating a final reconstruction at a global resolution of 2.95 Å. Since negligible structural difference at the catalytic active site was present between extended and non-extended forms, the two particle sets were combined to determine a structure with higher resolution at the core part of the polymerase. After a round of focused alignment with a mask including RNA and nsp12, 142,242 particles were finally selected and produced a final overall structure of Penciclovir-TP complex at a resolution of 2.60 Å by the same refinement procedures as above described.

For the samples of FAV-TP complex and RBV-TP complex, the steps of image processing including particle picking, 2D/3D classification and 3D refinement procedures were all performed using Relion 3.0. For FAV-TP complex, a total of 5,303,261 particles were automatically picked from 3,165 good micrographs. The extracted particles were subjected to a round of 2D classification and 3 rounds of 3D classification. Particle subsets from two high-resolution 3D reconstructions representing extended form and non-extended form of the nsp8, respectively, were merged and subjected to a round of focused alignment with a mask only including nsp12 to sort out particles with ordered conformation at the core part of the polymerase. After that, 166,089 particles were selected and produced a 2.91 Å 3D map. However, the bound Favipiravir at the active site showed poor density, indicative of the relatively low occupancy of polymerized ligand among the particle subset. Therefore, we performed an additional round of 3D classification with a small mask only encompassing the region around the +1 position of the primer strand. In this round of 3D classification, the alignment of rotations was disabled by using the *--skip_rotate* argument in Relion 3.0 and the Tau value was set to 80. One of the two subsequent classes corresponding to 116,511 particles had apparent strong density at the +1 position, whereas the other class clearly lacked the Favipiravir density. The final reconstruction of Favipiravir-TP complex was determined at a global resolution of 2.70 Å after CTF refinement and particle polishing.

For RBV-TP complex, 4,974,023 particles were automatically picked from 4,191 good micrographs. The extracted particles are subjected to a round of 2D classification, and a total of 1,077,748 particles were selected for two rounds of 3D classification and one additional round of focused alignment with a mask including nsp12. Finally, 196,511particles were sorted out and yielded a reconstruction of RBV-TP complex with a resolution of 2.97 Å after 3D refinement procedures as above described. The resolutions of all reconstructions were estimated based on the gold-standard Fourier shell correlation (FSC) = 0.143 criterion(*31*). The local resolution was calculated with Relion 3.0.

#### Model building

The model of the Remdesivir-ound RdRp complex of Covid-19 (PDB ID: 7BV2)(*9*) was used as the template for the model building of the three nucleotide-bound RdRp complexes. The template structure was docked into the density map using UCSF Chimera(*32*), followed by *ab initio* model building of the N-terminal NiRAN domain of nsp12 and one copy of nsp8, and manual adjustment and mutation of the ligands in COOT(*33*). Models were refined against corresponding density map using real_space_refine program in PHENIX(*34*). The model statistics were calculated with MolProbity(*35*) and listed in Table S1. Structural figures were prepared in Chimera or Chimerax(*36*).

#### Preparation of template-primer RNA for polymerase assays

The corresponding template-primer pair RNA strands were mixed at equal molar ratio in annealing buffer (10 mM Tris-HCl, pH 8.0, 25 mM NaCl and 2.5 mM EDTA). To anneal the RNA duplex, both oligonucleotides were denatured by heating to 94 °C for 5 min and then slowly cooled to room temperature. For the self-priming RNA2 used in the RBV bound RdRp complex, the RNA2 itself was denatured by heating to 94 °C for 5 min and then slowly cooled to room temperature.

#### RdRp enzymatic activity assay and its inhibition by nucleotide inhibitors

The purified SARS-CoV-2 RdRp complex from insect cell at final concentration of 1 μM was incubated with 3.0 μM dsRNA (Fig.1A) and 10 mM NTP (according to the corresponding dsRNA squence) in the presence of 1.14 U/μl RNase inhibitor in reaction buffer containing 20 mM Tris, pH8.0, 10 mM KCl, 6mM MgCl2, 0.01% Triton-X100, and 1mM DTT, which were prepared with DEPC-treated water. The total reaction volume was 20 μl. After incubation for 0 min, 5 min, 10 min, 20 min, 30 min, 45 min, and 60 min at a 37°C water bath, 40μl quench buffer (94% formamide, 30 mM EDTA, prepare with DEPC-treated water) was added to stop the reaction. A sample of 18 μl of reaction was mixed with 2 μl of 10x DNA loading buffer. Half of the sample (10 μl) was loaded onto a 10% or 20% Urea-PAGE denatured gel, run at 120V for 1h, and imaged with a Tanon-5200 Multi Fluorescence Imager.

The setup for the inhibition assays of the RdRp by the FAV-TP, RBV-TP, PEV-TP, favipiravir, ribavirin, and penciclovir is identical to the above for the RdRp enzymatic assays, except that FAV-TP, RBV-TP, PEV-TP, favipiravir, ribavirin, and penciclovir were added to final concentration of 0 μM, 1 μM, 10 μM, 100 μM, 1 mM, and 5 mM (also with an additional 10 mM in some cases) for 60 min before the addition of 10 mM NTP.

#### Synthesis of the triphosphate form of favipiravir (FAV-TP), ribavirin (RBV-TP) and penciclovir (PEV-TP)

For the synthesis of FAV-TP, The protective group was reacted at -2°C under perchloric acid. 1,2,3,5-tetraacetylribose reacted with favipiravir at 0°C for 24 hours, then sodium methoxide was added for deprotection, and quickly stirred for 5 minutes. And separated, synthesis intermediate. Favipiravir (100 mg) and trimethyl phosphate (1 mL) were added to a 100 mL three-necked flask, and replaced three times under vacuum-nitrogen conditions. Cool down to 0∼5°C in an ice-salt bath, and slowly add phosphorus oxychloride (30uL). The reaction system was stirred at 0°C for about 3 hours. Then add n-trioctylamine (190 mg), n-tributylamine pyrophosphate (290 mg) in N, N-dimethylformamide (3 mL) suspension. The reaction system was stirred at 4°C for 12 hours. The reaction solution was added with dichloromethane (25 mL) and stirred for 30 minutes, then filtered. After the filter cake was dissolved in water, the pH was adjusted to 7 with an aqueous sodium bicarbonate solution. The aqueous phase was extracted with methyl tert-butyl ether (5 mL*2) and concentrated at low temperature to remove water. The obtained residue was prepared and purified by HPLC and then lyophilized to obtain pure product.

For the synthesis of RBV-TP, Ribavirin (100 mg) and trimethyl phosphate (1 mL) were added to a 100 mL three-necked flask, and replaced three times under vacuum-nitrogen conditions. Cool down to 0∼5°C in an ice-salt bath, and slowly add phosphorus oxychloride (30uL). The reaction system was stirred at 0°C for about 3 hours. Then add n-trioctylamine (190 mg), n-tributylamine pyrophosphate (290 mg) in N,N-dimethylformamide (3 mL) suspension. The reaction system was stirred at 4°C for 12 hours. The reaction solution was added with dichloromethane (25 mL) and stirred for 30 minutes, then filtered. After the filter cake was dissolved in water, the pH was adjusted to 7 with an aqueous sodium bicarbonate solution. The aqueous phase was extracted with methyl tert-butyl ether (5 mL*2) and concentrated at low temperature to remove water. The obtained residue was prepared and purified by HPLC and then lyophilized to obtain pure product.

For the synthesis of PEV-TP, add penciclovir (100 mg) and trimethyl phosphate (1 mL) to a 100 mL three-necked flask, and replace with vacuum-nitrogen three times. Cool down to 0∼5°C in an ice-salt bath, and slowly add phosphorus oxychloride (30uL). The reaction system was stirred at 0°C for 36 hours and quenched to purify the pure penciclovir monophosphate required for one side. Then add n-trioctylamine (190 mg), n-tributylamine pyrophosphate (290 mg) in N,N-dimethylformamide (3 mL) suspension. The reaction system was stirred at 0°C for 12 hours.

The reaction solution was added with dichloromethane (25 mL) and stirred for 30 minutes, then filtered. After the filter cake was dissolved in water, the pH was adjusted to 7 with sodium bicarbonate aqueous solution. The aqueous phase was extracted with methyl tert-butyl ether (5 mL*2) and concentrated at low temperature to remove water. The obtained residue was prepared by HPLC and lyophilized to obtain pure product.

#### Pharmacokinetic Studies

The PK studies were conducted at Suzhou HQ Bioscience Co., Ltd. The animals (N = 3 for SD rats) were fasted for 12 h before the administration of tested compounds. For PK studies in rats, each compound dissolved in DMSO-enthanol-PEG300-saline (5/5/40/50, v/v/v/v) was administered orally at 10 mg/Kg or intravenously at 2 mg/Kg, respectively. Blood samples were collected from the jugular vein into EDTA-K2 tubes at various time points post-dose. Serum samples were obtained following general procedures and the concentrations of analytes in the supernatant were analyzed by LC-MS/MS.

#### Tissue Distribution Study of the metabolites of X3 in mice

Sixteen male CD-1 mice were randomly divided into four groups (n = 4, each). All mice were fasted for 12 h before administered orally with a single dose of 200 mg/kg X3 dissolved in DMSO-enthanol-PEG300-saline (5/5/40/50, v/v/v/v). At 1, 2, 4, and 8 h post-dosing, the mice were anesthetized, and tissues including heart, liver, lung, kidney and testis were harvested. Tissue samples were individually homogenized, and the concentrations of X1, X1-MP and X1-TP in homogenates were analyzed by LC-MS/MS.

#### Cell lines and viruses

African green monkey kidney Vero E6 cells (ATCC-1586) were maintained in Dulbecco’s modified Eagle’s medium (DMEM) with 10% fetal bovine serum (FBS) and 1% penicillin– streptomycin antibiotics. Cells were kept at 37 °C in a 5% CO_2_ atmosphere. The strain (nCoV-2019BetaCoV/Wuhan/WIV04/2019) of SARS-CoV-2, which was isolated from a clinical patient, was obtained from National Virus Resource Center. The virus strain was propagated in Vero E6 cells.

#### Antiviral activities and Cytotoxicity measurement for nucleotide analogs

In our study, Vero E6 cells were pre-seeded to 48-well plates (50,000 cells/well) for 16-18 h, and treated with medium containing gradient concentration of nucleotide analogs at 100 μL/well for 1 h. Then, the cells were inoculated with SARS-CoV-2 at multiplicity of infection (MOI) of 0.05. One hour later, the supernatant was removed, and cells were washed with PBS, and treated with fresh medium containing gradient concentration of nucleotide analogs at 200 μL/well. At 24 h post infection, the cell supernatant was collected, antiviral activities were evaluated by quantification of viral copy numbers in the cell supernatant via real-time fluorescence quantitative PCR (qRT-PCR)(TaKaRa MiniBEST Viral RNA/DNA Extraction Kit Ver.5.0 TaKara 9766,PrimeScript™ RT reagent Kit with gDNA Eraser TaKaRa RR047A, TaKaRa SYBR® Premix Ex Taq ™ II TaKaRa RR820A. The inhibition rate of nucleotide analogs was calculated based on the viral copy number, and the 50% maximal effective concentration (EC_50_) was calculated with Graphpad Prism software 8.0. The experiments were done in triplicates and all the infection experiments were performed at biosafety level 3 (BSL-3).

For cytotoxicity measurement, Vero E6 cells were added to a 96-well plate (20,000 cells/well), and added with medium containing gradient concentration of nucleotide analogs at 100 μL/well next day. The 50% cytotoxicity of VV116 was determined after 24 h using the CCK8 assay kit, and CC_50_ of nucleotide analogs was calculated with Graphpad Prism software 8.0.

#### Mice

The BALB/c mice were bred and maintained in specific pathogen free (SPF) environment at the Laboratory Animal Center of Wuhan Institute of Virology, CAS. Eleven female mice at six to eight weeks-old were used in our study. All virus infection experiments are conducted in biosafety Level 3 (BSL3) facility. All of our animal experiments were conformed to the use and care of laboratory animals and the Institutional Review Board of the Wuhan Institute of Virology, CAS.

The experiment was divided into two groups, the Ad5-hACE2-SARS-CoV-2-VV116 (Ad5-VV116 for short) and the Ad5-hACE2-SARS-CoV-2-Vehicle (Ad5-Vehicle for short) group. Mice were anesthetized by intraperitoneal injection of 2.5% Avertin (20 μL/g body weight) and transduced intranasally with 50 μL 5×10^9^ PFU/ml of Ad5-ACE2. One day prior to infection SARS-CoV-2, Ad5-VV116 treatment with VV116 and continued dosing of 100 mg/kg once daily until the mice were sacrificed at day 5, Ad5-Vehicle received normal saline as control. At day 0, mice were anesthetized as above and infected with 50 μL 2×10^6^ PFU/ml of SARS-CoV-2. Mice were lightly anesthetized with isoflurane and weighted and observed for clinical signs daily from day 0 to day 5.

Mice were sacrificed at day 5 after intranasally injection of SARS-CoV-2. Left Lung were fixed with 4% paraformaldehyde, Fixed tissue samples were used for hematoxylin-eosin (H&E) staining and Immunohistochemistry (IHC) for the detection of the hACE2 (R&D, 171606; 1:1000 dilution) and immunofluorescence (IFA) for the detection of the SARS-CoV-2 NP (the NP of a bat SARS-related CoV; 1:500 dilution) antigen. Right lung tissues from euthanized mice were homogenized with DMEM, the homogenized tissues were centrifuged at 3000 rpm for 10 min at 4 °C, and RNA was extracted from the supernatant by RNeasy Mini Kit (Qiagen, 74104) for real-time quantitative PCR (RT-PCR) testing.

#### Synthesis of compounds X1, X2, and X3

The materials and reagents were commercially available and solvents, if necessary, were purified and dried by standard methods. ^1^H-NMR and ^13^C-NMR spectra were determined on a Brucker 400 Hz, Brucker 500 Hz or Brucker 600 Hz instrument. ESI-MS was determined on Finnigan™ LTQ™ (*Thermo Fisher Scientific*, Bremen, Germany) linear ion trap mass spectrometer. All reactions were monitored by thin-layer chromatography (TLC) on 25.4×76.2 mm silica gel plates (GF-254). The final compounds possessed a HPLC purity of ≥ 95%.

#### Preparation of compound X1

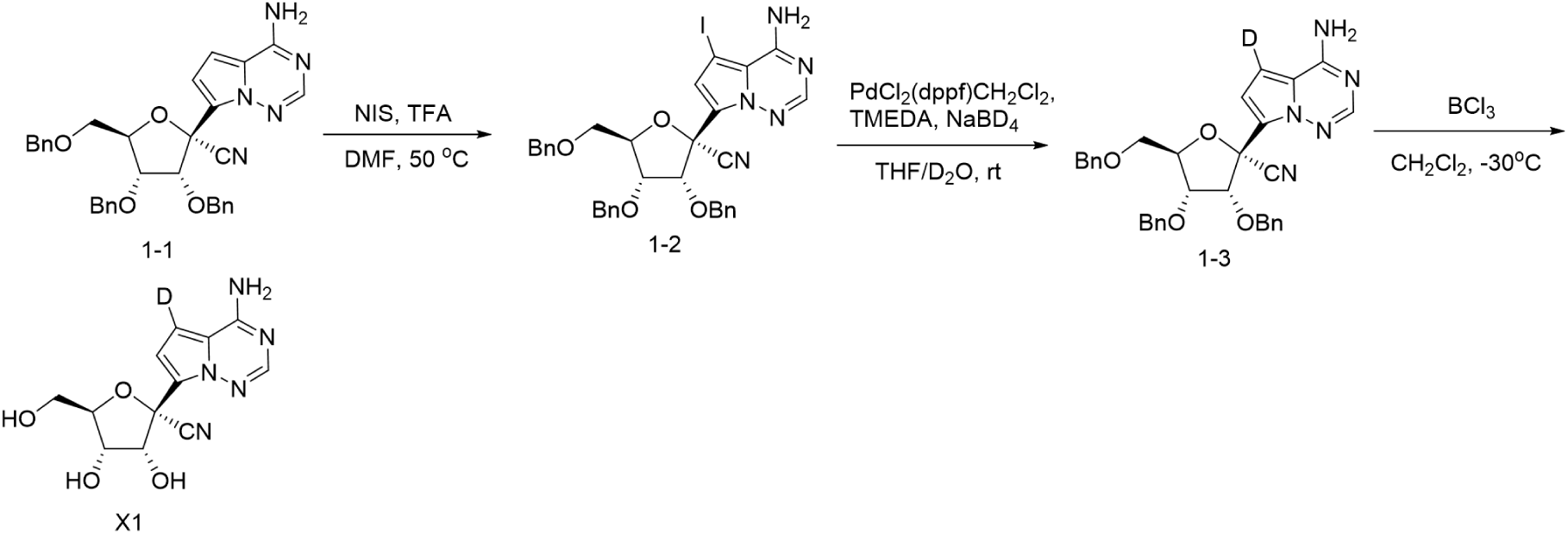

Compound **1-1** was prepared according to the previously reported method (Nature 2016, 531, 381). To a solution of **1-1** (20 g, 35.6 mmol) in DMF (70 mL) was added NIS (8.8 g, 39.2 mmol) and CF_3_COOH (0.81 g, 7.1 mmol). The mixture was stirred at 50 °C under N_2_ protection for 1 h, then poured into a solution of Na_2_SO_3_ (9.0 g,71.2 mmol) and Na_2_CO_3_ (15.1 g,142.4 mmol) in water (150 mL), and extracted with ethyl acetate. The exact was washed with brine, dried over Na_2_SO_4_, and concentrated under vacuum. The resulting crude product was slurried in isopropyl ether and filtered to give **1-2** as a white solid (21.5 g, yield 88%). ^1^H NMR (400 MHz, CDCl_3_) δ 7.86 (s, 1H), 7.42 – 7.16 (m, 15H), 7.01 (s, 1H), 6.22 (s, 2H), 4.94 – 4.85 (m, 2H), 4.71 (d, *J* = 5.0 Hz, 1H), 4.62 – 4.52 (m, 2H), 4.51 – 4.41 (m, 2H), 4.32 (d, *J* = 12.0 Hz, 1H), 4.03 (dd, *J* = 7.0, 4.9 Hz, 1H), 3.82 (dd, *J* = 11.0, 3.3 Hz, 1H), 3.62 (dd, *J* = 11.0, 3.7 Hz, 1H).

Compound **1-2** (1.77 g, 2.57 mmol) was added to anhydrous tetrahydrofuran (20 mL) and deuteroxide (2 mL), and then the solvent was removed in vacuum. The procedure was repeated, followed by addition of tetrahydrofuran (20 mL) and deuteroxide (2 mL) again. To this solution were added 1,1’-bis(diphenylphosphino)ferrocene-palladium(II)dichloride dichloromethane complex (0.11 g, 0.13 mmol) and N,N,N’,N’-tetramethylethylenediamine (0.60 g, 5.14 mmol). After stirring for 10 min at room temperature, sodium borodeuteride (0.22 g, 5.14 mmol) was added in portion within 30 min. The mixture continued to be stirred at room temperature until the starting material disappeared completely. Then, the reaction was quenched with saturated NH_4_Cl solution, and extracted with ethyl acetate. The organic phase was washed with brine, dried over Na_2_SO_4_, and concentrated. The residue was purified by chromatography on silica gel to give compound **1-3** with a deuterium content > 98% as a white solid (0.79 g, yield 55%).

Debenzylation of compound **1-3** (0.56 g, 1.0 mmol) with 1M BCl_3_ dichloromethane solution (3.5 mmol, 3.5 mL) was performed according to the existing method (Nature 2016, 531, 381) to afford compound **X1** as a white solid (0.15 g, yield 53%). ^1^H NMR (500 MHz, DMSO-*d*_*6*_) δ 8.01 – 7.79 (m, 3H), 6.88 (s, 1H), 6.09 (d, *J* = 6.4 Hz, 1H), 5.19 (d, *J* = 5.3 Hz, 1H), 4.91 (t, *J* = 5.8 Hz, 1H), 4.65 (t, *J* = 5.7 Hz, 1H), 4.08 – 4.04 (m, 1H), 3.98 – 3.93 (m, 1H), 3.67 – 3.61 (m, 1H), 3.54 – 3.48 (m, 1H). ^13^C NMR (126 MHz, DMSO-*d*_6_) δ 156.08, 148.32, 124.35, 117.82, 116.94, 111.16, 85.92, 79.04, 74.72, 70.56, 61.43. MS m/z = 293.0 [M + 1]^+^.

#### Preparation of compound X2

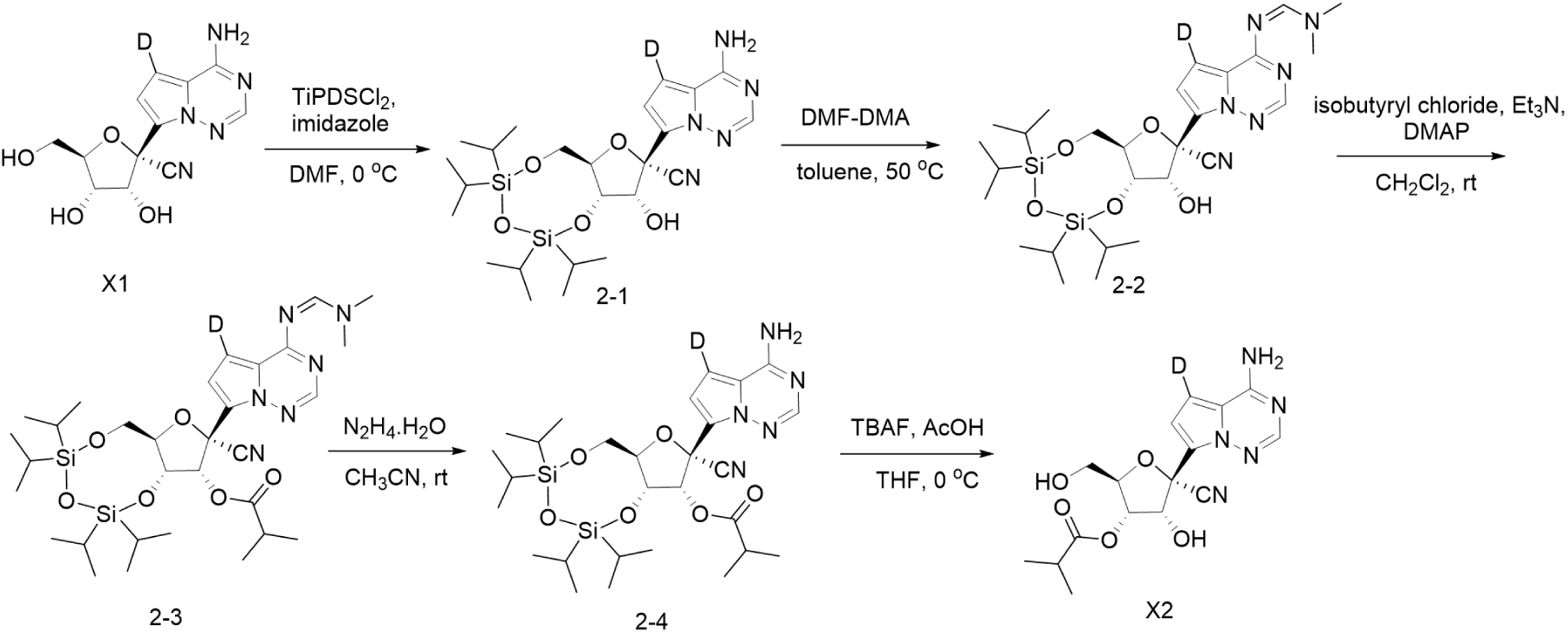

To a solution of compound **X1** (879 mg, 3.01 mmol) and imidazole (819 mg, 12.03 mmol) in DMF (15 mL) was added 1,3-dichloro-1,1,3,3-tetraisopropyldisiloxane (1.32 g, 4.21 mmol) dropwise under ice bath. After the addition, the mixture was stirred at room temperature for 4 h, then poured into water (100 mL), and extracted with ethyl acetate. The organic layer was separated, washed with brine, dried over Na_2_SO_4_ and concentrated. The residue was added in petroleum ether, and stirred overnight. The resulting solid was collected by filtration and dried to give compound **2-1** as a white solid (1.29 g, yield 80%). ^1^H NMR (600 MHz, DMSO-*d*_*6*_) δ 8.04 – 7.78 (m, 3H), 6.79 (s, 1H), 6.46 (d, *J* = 5.7 Hz, 1H), 4.56 (t, *J* = 5.1 Hz, 1H), 4.24 – 4.08 (m, 3H), 3.91 (d, *J* = 12.0 Hz, 1H), 1.08 – 0.86 (m, 28H).

Compound **2-1** (1.10 g, 2.06 mmol) was added in toluene (20 mL) followed by the addition of N-dimethylformamide dimethyl acetal (370 mg, 3.11 mmol). The mixture was stirred at 45 °C for about 30 min. The solvent was removed under vacuum to give compound **2-2** as a white foam (1.12g, yield 92%).

To a solution of compound **2-2** (614 mg, 1.04 mmol) in dichloromethane (10 mL) were successively added triethylamine (210 mg, 2.08 mmol), isobutyryl chloride (166 mg, 1.56 mmol) and DMAP (127 mg, 1.04 mmol). The mixture was stirred at room temperature for 1 h, and then treated with saturated sodium bicarbonate solution and dichloromethane. the organic layer was separated, dried over Na_2_SO_4_, concentrated, and purified by chromatography on silica gel to give compound **2-3** as a white foam (494 mg, yield 72%).

Compound **2-3** (350 mg, 0.53 mmol) was added in acetonitrile (8 mL) followed by the addition of 85% hydrazine hydrate (125 mg, 2.12 mmol) at room temperature. The mixture was stirred for about 30 min, then poured into water, and extracted with ethyl acetate. The organic layer was separated, washed with 1M hydrochloric acid, saturated sodium bicarbonate and brine, then dried over Na_2_SO_4_, and evaporated to give compound **2-4** as a white solid (289 mg, yield 90%). To a solution of compound **2-4** (289 mg, 0.48 mmol) and acetic acid (7 mg, 0.12 mmol) in tetrahydrofuran (10 mL) was added 1 M solution of tetrabutylammonium fluoride in THF (0.48 mL, 0.48 mmol) under ice bath. The mixture was stirred for 2-3 h, then poured into water, and extracted with isopropyl acetate. The organic layer was separated, washed with saturated sodium bicarbonate and brine, then dried over Na_2_SO_4_ and evaporated. The crude product was slurried in n-heptane/isopropanol, and the resulting precipitate was filtrated to give compound **X2** as a white solid (121 mg, yield 70%). ^1^H NMR (600 MHz, DMSO-*d*_*6*_) δ 8.06 – 7.85 (m, 3H), 6.88 (s, 1H), 6.41 (d, *J* = 6.5 Hz, 1H), 5.21 (dd, *J* = 5.7, 3.3 Hz, 1H), 5.08 – 5.04 (m, 1H), 4.99 (t, *J* = 6.0 Hz, 1H), 4.26 (q, *J* = 3.7 Hz, 1H), 3.64 – 3.52 (m, 2H), 2.67 – 2.58 (m, 1H), 1.17 (d, *J* = 7.0 Hz, 3H), 1.15 (d, *J* = 7.0 Hz, 3H). ^13^C NMR (126 MHz, DMSO-*d*_6_) δ 175.92, 156.13, 148.45, 123.24, 117.50, 117.27, 111.60, 84.59, 78.30, 73.14, 72.69, 61.25, 33.81, 19.18, 19.08. MS m/z = 363.0 [M + 1]^+^. MS m/z = 363.0 [M + 1]^+^.

### Preparation of compound X3

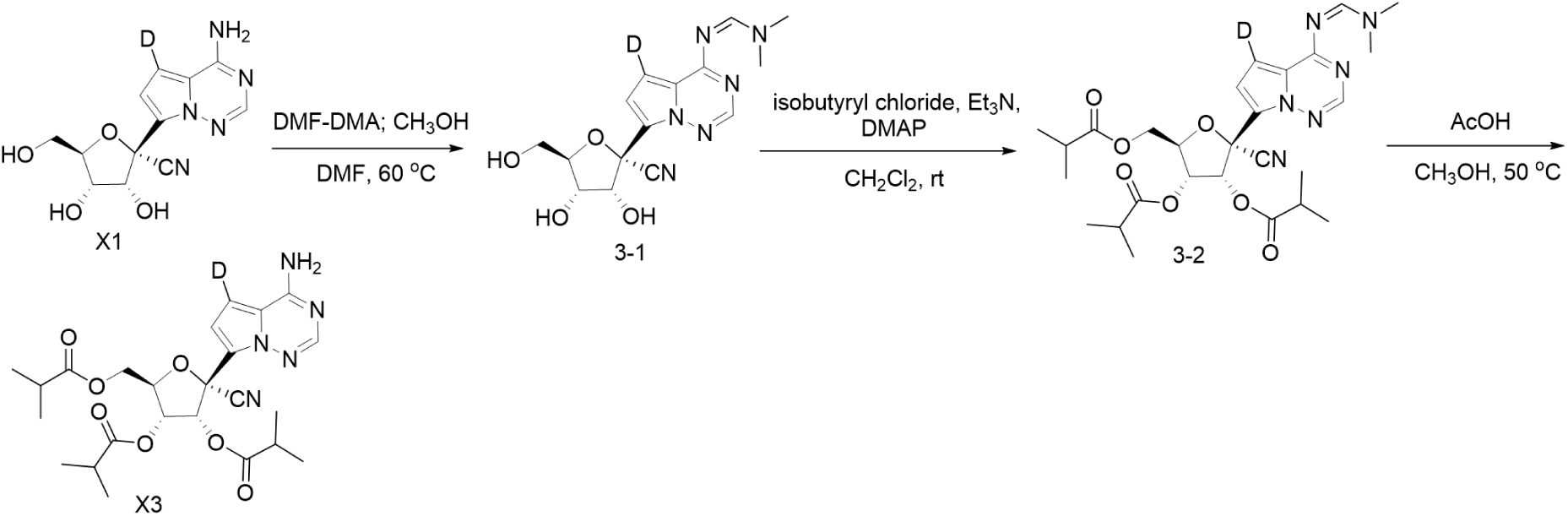

To a solution of compound **X1** (0.58 g, 2.0 mmol) in DMF (10 mL) was added N,N-dimethylformamide dimethyl acetal (1.43 g, 12.0 mmol) at room temperature. The mixture was stirred at 60 °C for 2 h, then cooled to room temperature, and an equivalent volume of methanol was added. The solvent was evaporated under vacuum to give an oily residue which was treated with isopropyl alcohol. The resulting precipitate was collected by filtration and dried to give compound **3-1** as an off-white solid (0.55 g, 80% yield).

Compound **3-1** (0.37 g, 1.0 mmol), triethylamine (0.81 g, 8.0 mmol) and DMAP (0.37 g, 3.0 mmol) were suspended in methylene chloride (10 mL), and to this solution was slowly added isobutyryl chloride (0.43 g, 4.0 mmol) at room temperature. The mixture was stirred until the starting material completely disappeared. The solvent was removed under vacuum, and the residue was partitioned between water and ethyl acetate. The organic layer was separated, washed successively with 1M hydrochloric acid, saturated sodium bicarbonate and brine, then dried over Na_2_SO_4_, and evaporated to give the crude product **3-2** as an oil, which was taken directly to the following step without further purification.

The obtained intermediate **3-2** was dissolved in ethanol (10 mL), followed by the addition of acetic acid (1.2 g, 20.0 mmol) at room temperature. The mixture was stirred at 50 °C overnight, and concentrated. The resulting residue was purified by chromatography on silica gel to afford compound **X3** as a white foam (0.38 g, 75% yield for two steps). ^1^H NMR (500 MHz, DMSO-*d*_6_) δ 8.04 (br, 1H), 7.98 (br, 1H), 7.94 (s, 1H), 6.77 (s, 1H), 6.09 (d, *J* = 5.7 Hz, 1H), 5.45 (dd, *J* = 5.8, 3.7 Hz, 1H), 4.64 (q, *J* = 3.7 Hz, 1H), 4.34 (dd, *J* = 12.4, 3.3 Hz, 1H), 4.29 (dd, *J* = 12.4, 4.1 Hz, 1H), 2.69 – 2.56 (m, 2H), 2.51 – 2.46 (m, 1H), 1.19 (d, *J* = 7.1 Hz, 3H), 1.16 (d, *J* = 6.9 Hz, 3H), 1.14 – 1.10 (m, 6H), 1.06 (d, *J* = 7.0 Hz, 3H), 1.03 (d, *J* = 7.0 Hz, 3H). ^13^C NMR (126 MHz, DMSO-*d*_6_) δ 175.53, 174.90, 174.13, 155.58, 148.12, 120.98, 117.17, 115.44, 110.30, 81.25, 75.81, 72.05, 70.30, 62.46, 33.20, 33.16, 33.09, 18.55, 18.46, 18.40, 18.35, 18.33, 18.18. MS m/z = 503.0 [M + 1]^+^.

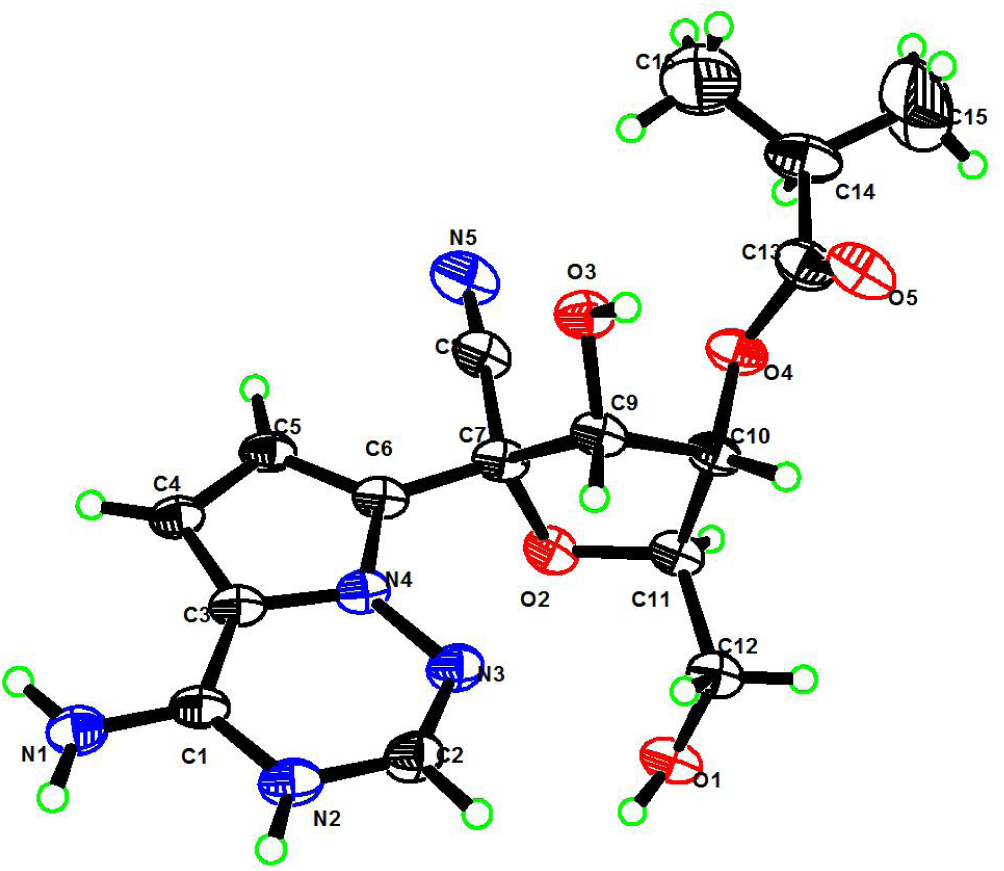

The X-ray crystal structure of **X2**

^1^H NMR and ^13^C NMR spectrum of X1, X2 and X3.

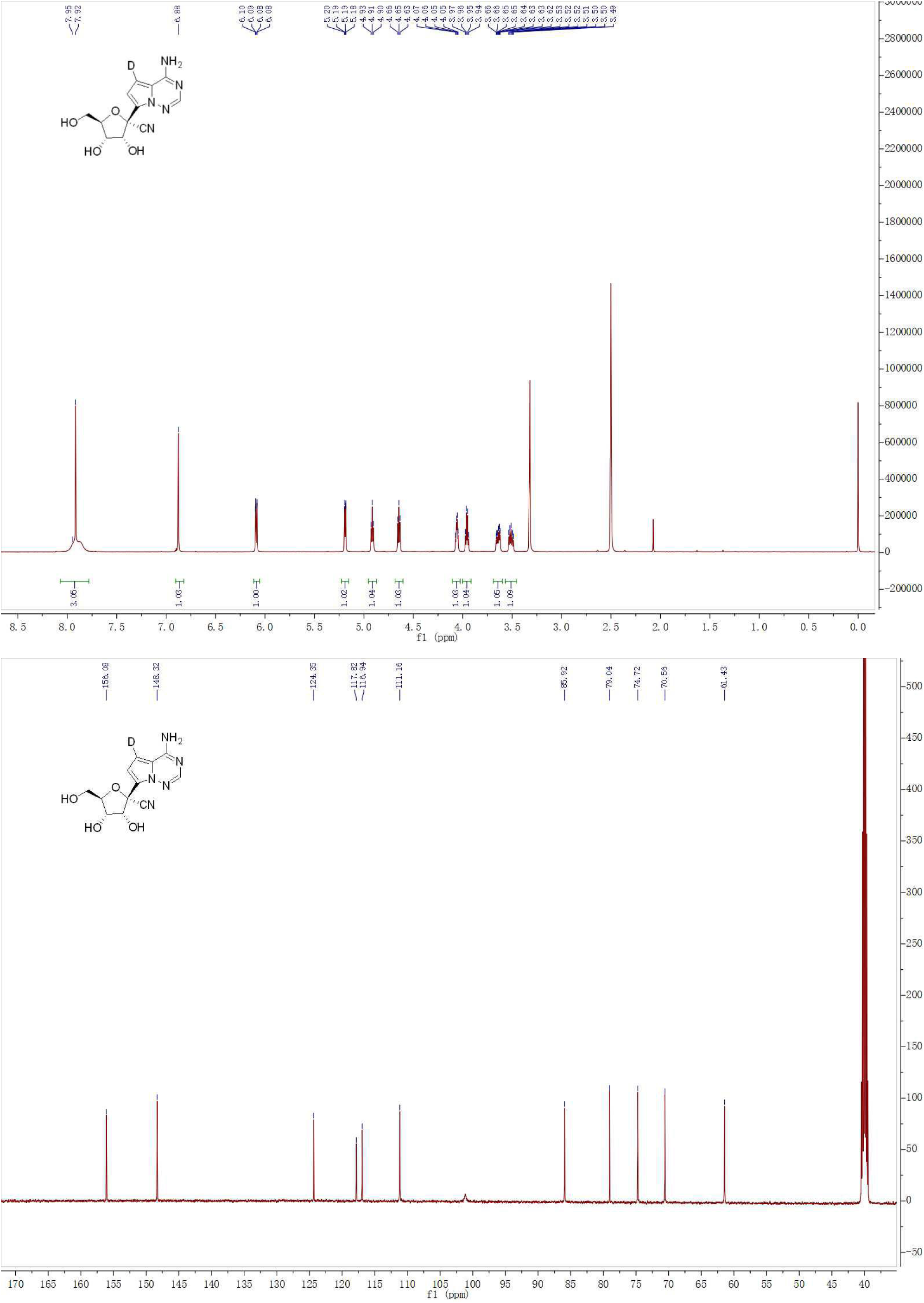

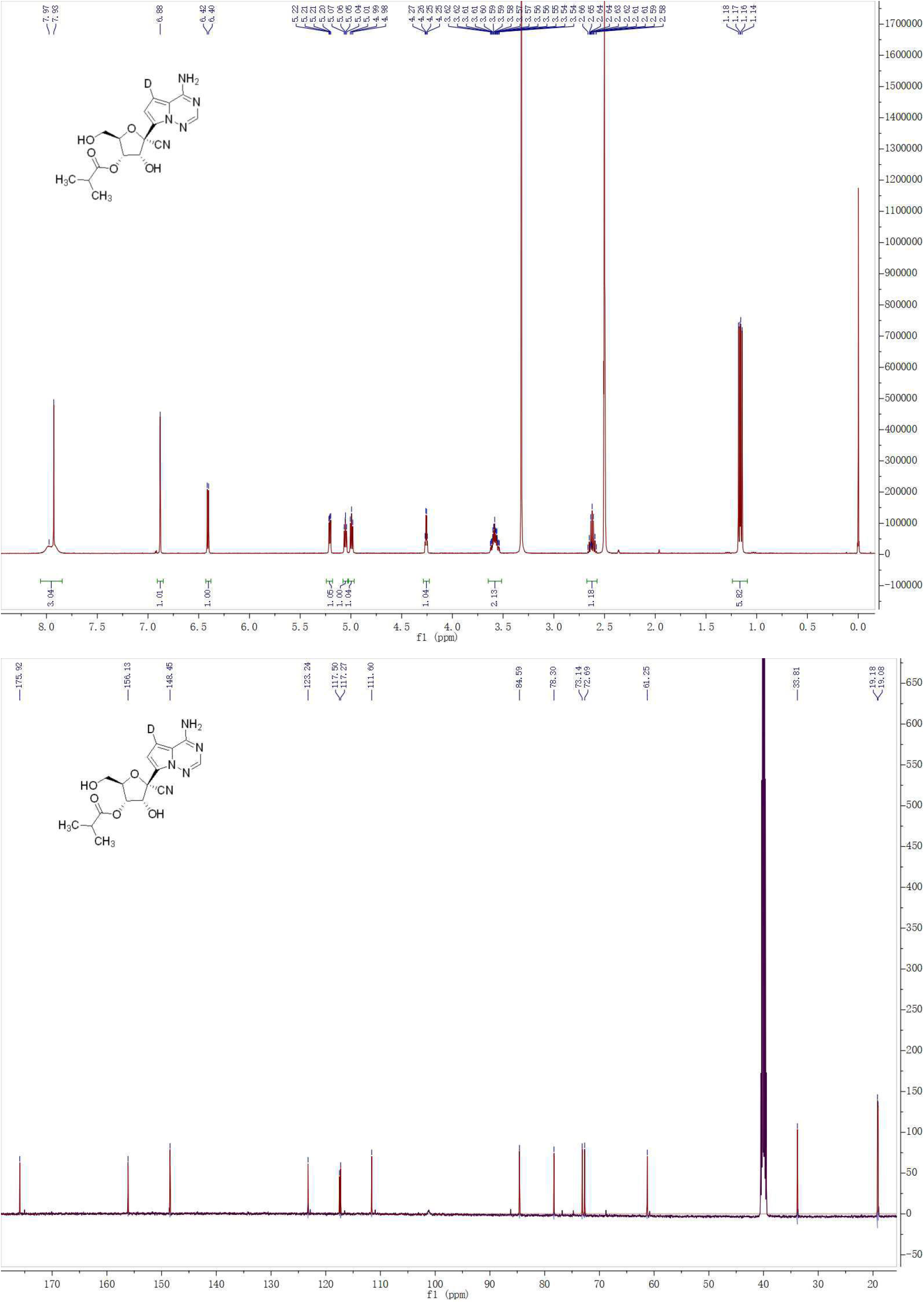

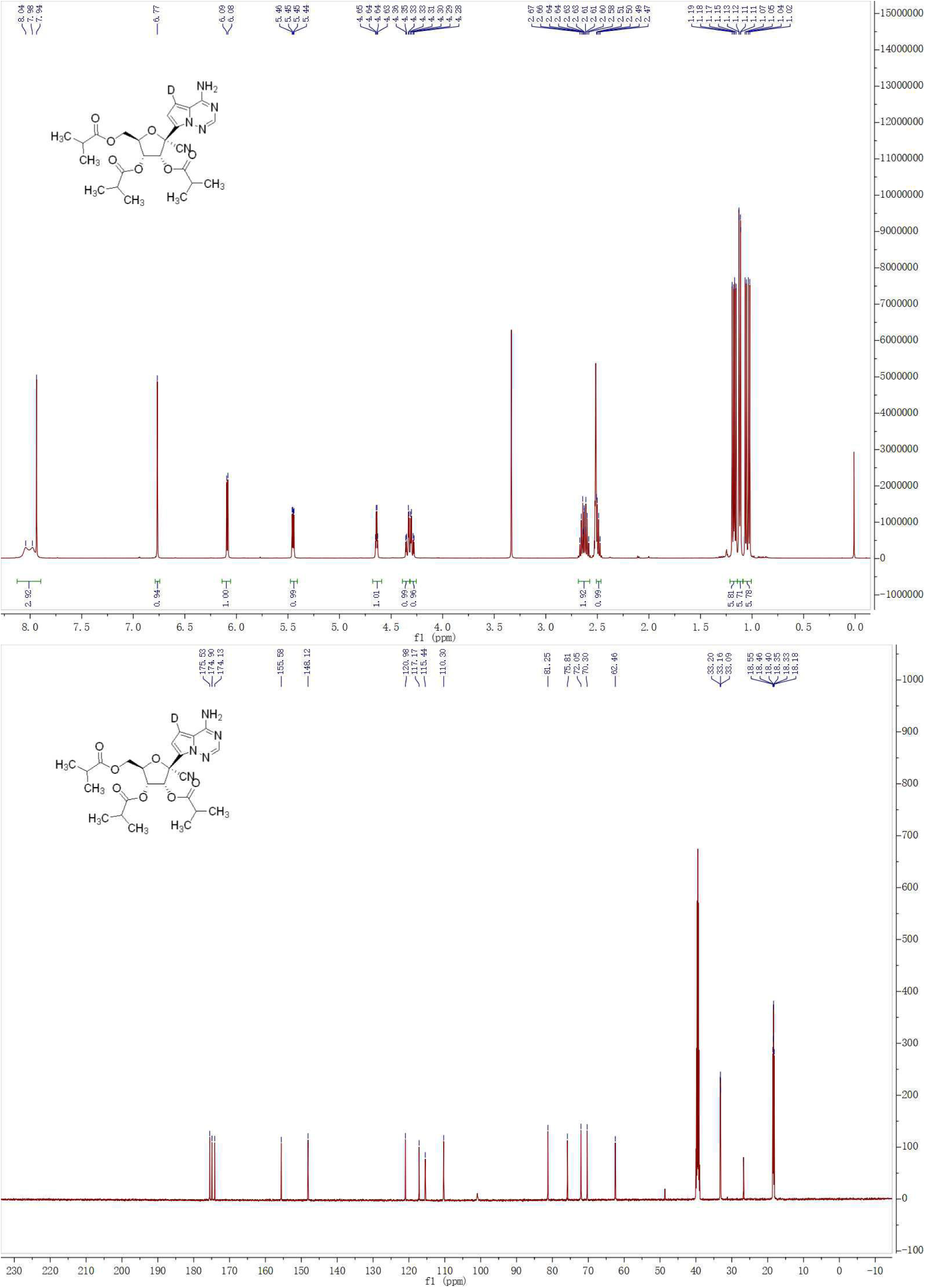

**Table S1.** Cryo-EM data collection, refinement, and validation statistics.

|  | RBV | FAV | PEV-overall | PEV-extended |
| --- | --- | --- | --- | --- |
| **Data collection and Processing (for each dataset)** |  |  |  |  |
| Microscope | Titan Krios | Titan Krios | Titan Krios | Titan Krios |
| Voltage (keV) | 300 | 300 | 300 | 300 |
| Camera | Gatan K3™ | Gatan K3™ | Gatan K3™ | Gatan K3™ |
| Magnification | 46773 | 46773 | 46773 | 46773 |
| Pixel size at detector (Å/pixel) | 1.069 | 1.069 | 1.069 | 1.069 |
| Total electron exposure (e-/Å<sup>2</sup>) | 69 | 69 | 69 | 69 |
| Exposure rate (e-/pixel/sec) | 26 | 26 | 26 | 26 |
| Number of frames collected during exposure | 36 | 36 | 36 | 36 |
| Defocus range (µm) | -0.7 to -2.6 | -0.4 to -2.5 | -0.4 to -2.3 | -0.4 to -2.3 |
| Phase plate (if used) | N/A | N/A | N/A | N/A |
| -phase shift range (in degrees) | N/A | N/A | N/A | N/A |
| -number of images per phase plate position | N/A | N/A | N/A | N/A |
| Automation software (EPU, SerialEM or manual) | SerialEM | SerialEM | SerialEM | SerialEM |
| Tilt angle (if grid was tilted) | 0 | 0 | 0 | 0 |
| Energy filter slit width (if used) | 20 | 20 | 20 | 20 |
| Micrographs collected (no.) | 4297 | 3209 | 3913 | 3913 |
| Micrographs used (no.) | 4191 | 3165 | 3893 | 3893 |
| Total extracted particles (no.) | 4974023 | 5303261 | 5240500 | 5240500 |
| **For each reconstruction:** |  |  |  |  |
| Refined particles (no.) | 199679 | 116511 | 142242 | 139773 |
| Final particles (no.) | 199679 | 116511 | 142242 | 139773 |
| Point-group or helical symmetry parameters | C1 | C1 | C1 | C1 |
| Resolution (global, Å) |  |  |  |  |
| FSC 0.5 (unmasked/masked) | 4.4/3.3 | 4.3/3.0 | 4.4/3.0 | 4.6/3.7 |
| FSC 0.143 (unmasked/masked) | 3.8/3.0 | 3.5/2.7 | 3.6/2.6 | 3.1/2.7 |
| Resolution range (local, Å) | 2.9-6.0 | 2.5-6.0 | 2.5-6.0 | 2.5-6.0 |
| Map sharpening B factor (Å<sup>2</sup>) | -69 | -46 | -32 | -45 |
| **Model composition (for each model)** |  |  |  |  |
| Protein | 1191 | 1200 | 1195 | 1323 |
| Ligands | 8 | 9 | 9 | 9 |
| RNA/DNA | 20 | 28 | 27 | 44 |
| **Model Refinement (for each model)** |  |  |  |  |
| Refinement package | PHENIX-1.14_3260 | PHENIX-1.14_3260 | PHENIX-1.14_3260 | PHENIX-1.14_3260 |
| -real or reciprocal space | real space | real space | real space | real space |
| Initial model used (PDB code) | 7BV2 | 7BV2 | 7BV2 | 7BV2 |
| Model-Map CC | 0.82 | 0.87 | 0.85 | 0.85 |
| Model resolution (Å) | 3.2 | 2.8 | 2.8 | 2.9 |
| FSC threshold | 0.5 | 0.5 | 0.5 | 0.5 |
| B factors (Å<sup>2</sup>) |  |  |  |  |
| Protein residues | 68.01 | 51.68 | 49.7 | 52.23 |
| Ligands | 49.28 | 28.14 | 30.04 | 27.16 |
| RNA/DNA | 97.28 | 98.03 | 82.32 | 117.15 |
| R.m.s. deviations from ideal values |  |  |  |  |
| Bond lengths (Å) | 0.009 | 0.012 | 0.015 | 0.008 |
| Bond angles (°) | 0.935 | 0.965 | 1.055 | 1.016 |
| **Validation (for each model)** |  |  |  |  |
| MolProbity score | 1.52 | 1.34 | 1.46 | 1.38 |
| CaBLAM outliers | 1.89 | 0.94 | 1.54 | 1.23 |
| Clashscore | 5.64 | 4.22 | 5.73 | 5.63 |
| Poor rotamers (%) | 0.09 | 0.38 | 0.19 | 0.6 |
| C-beta deviations | 0.00 | 0.00 | 0.00 | 0.00 |
| EMRinger score (if better than 4 Å resolution) | 2.92 | 4.18 | 4.42 | 3.23 |
| Ramachandran plot |  |  |  |  |
| Favored (%) | 96.6 | 97.3 | 97.21 | 97.64 |
| Outliers (%) | 0.00 | 0.00 | 0.00 | 0.00 |

